# Bioelectrical phase transitions

**DOI:** 10.64898/2026.07.07.734602

**Authors:** Joshua B. Fernandes, Hyeongjoo Row, Karthik Shekhar, Kranthi K. Mandadapu

## Abstract

Electrical signaling in biological systems is generally understood through the lens of single-channel biophysics, yet whether ensembles of ion channels can undergo cooperative opening and closing remains unclear. Here, we show that ensembles of voltage-gated ion channels can undergo bioelectrical order–disorder phase transitions driven by feedback between channel currents and local membrane voltage. When channels open, they carry ion-selective current that redistributes ions near the membrane and perturbs the transmembrane potential, thereby biasing the gating of nearby channels. This emergent nonequilibrium coupling generates a bona fide phase transition in ion channel ensembles. Finite-size analyses of the open-channel fraction, its fluctuations, and the distribution of collective channel states yield a voltage–temperature phase diagram with a first-order line separating collectively open and closed states and terminating at a critical point. The critical temperature is governed by a dimensionless conductance ratio set by ion transport, channel density, and confinement geometry. Applying this framework to measurements from the squid giant axon, the axon initial segment, and the nodes of Ranvier suggests that collective activation may be favored by high sodium-channel densities in large-diameter nerves, whereas the lower densities typical of potassium channels place them in an independent-gating regime.

## Introduction

The fundamental basis of electrical signaling rests on the voltage-dependent responses of single ion channels and their associated current–voltage relationships [1, 2]. These single-channel characteristics underpin the modern understanding of neuronal signaling phenomena, including action potential initiation and propagation by voltage-gated Na^+^ and K^+^ channels, and presynaptic neurotransmitter release triggered by voltage-gated Ca^2+^ channels [2, 3]. However, many biological systems, notably axonal structures [4, 5] and a broad class of excitable cells [3, 6], contain high densities of ion channels whose collective behavior may shape macroscopic electrophysiological responses. In such dense assemblies, an open channel does not merely respond to the local voltage; by carrying ion-selective current, it can also perturb the voltage experienced by nearby channels. While the properties of individual channels have been characterized in considerable detail, the emergent behavior of collections of channels remains poorly understood. In particular, it is an open question whether cooperative gating, i.e., correlated collective opening and closing of channel ensembles under an applied voltage, can arise in such systems.

In this work, starting from single-channel behavior, we show that a model of an ensemble of ion channels exhibits cooperative behavior consistent with a thermodynamic phase transition (Fig. 1). Here, a phase transition refers to a collective switch in the state of the ensemble: many channels change state together, so that a small change in voltage or temperature produces a sharp macroscopic change in the fraction of open channels. Specifically, we find a first-order transition line separating collectively open and closed states in the voltage–temperature plane, terminating at a critical point, as schematized in Fig. 1b. We further show that the critical temperature is controlled by a dimensionless conductance ratio, providing a criterion for when collective activation may occur in biological or reconstituted membrane systems. Such phase-transition behavior adds to the growing catalog of emergent phenomena identified in biological systems, including the condensation transitions that govern the organization of the cytoplasm and other intracellular compartments [7, 8].

**Figure 1:**
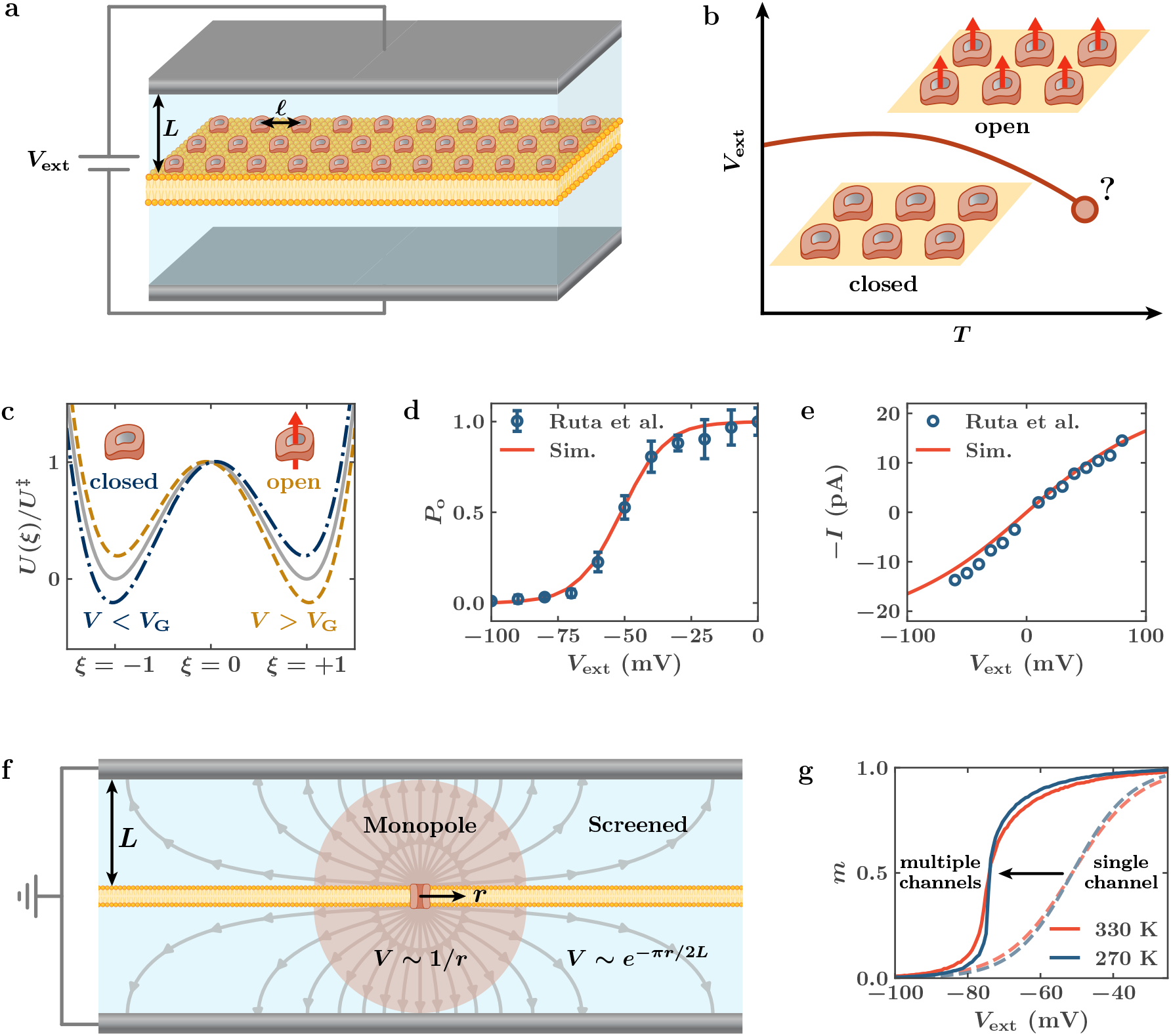
From single-channel behavior to multichannel cooperativity. (a) Schematic of the model system consisting of a large flat membrane containing voltage-gated K^+^ channels arranged on a square lattice with spacing *ℓ*, placed between two plate electrodes a distance *L* from the membrane and maintained at an applied potential difference *V*_ext_. (b) Schematic of a phase transition between collectively closed and collectively open states in the voltage–temperature plane, with a first-order line terminating at a critical point. (c) Channel gating is modeled by a single state variable *ξ* evolving in a double-well potential with two minima, where *ξ* < 0 corresponds to a closed channel state and *ξ* > 0 corresponds to an open channel state, separated by an energy barrier *U* ^‡^. The free-energy landscape in *ξ* is biased by the local transmembrane voltage *V* − *V*_G_, with the closed and open states equiprobable when *V* = *V*_G_. (d,e) Simulations of a single KvAP channel fit the long-time open probability (d) and open-channel current (e) measured by Ruta et al. [9], leading to the parameter values *V*_G_ = −50 mV, *Q* = 1.6*e, R*_P_ = 0.7 nm, and *k* = 350 pS. (f) Current through an ion-selective open channel produces a long-ranged monopolar perturbation to the membrane potential within a distance *r* < *L* and an exponentially screened perturbation for *r* > *L*, which we neglect. (g) Simulations of the coupled dynamics of 56 × 56 channels with spacing *ℓ* = 3 nm and confinement length *L* = 27 nm, with all other parameters as in panels (d,e), show a shift of approximately −25 mV in the threshold for collective activation compared to an isolated channel, indicating emergent cooperativity. The isolated-channel activation curves are the same as in panel (d).

### Single channel kinetics

We begin with the single-channel behavior that forms the microscopic basis for the emergence of collective behavior. As shown schematically in Fig. 1a, we consider a membrane containing voltage-gated ion channels immersed in a symmetric binary electrolyte and subjected to an externally applied voltage *V*_ext_, with electrodes located a distance *L* from the membrane. We focus primarily on the prokaryotic KvAP potassium channel as a model system, in part because its single-channel electrophysiological properties, including voltage gating and current–voltage characteristics, have been measured in detail [9]. Moreover, KvAP shares key structural and functional features with eukaryotic Kv channels, particularly in the selectivity filter and voltage-dependent gating machinery [9, 10]. Thus, KvAP provides a useful calibrated example in which the single-channel gating and current can be constrained by experiment. While KvAP provides a calibrated model system, the mechanism identified here relies only on voltage-dependent gating and ion-selective current through the membrane. We therefore expect the collective behaviors described here to apply more broadly to ensembles of voltage-gated ion channels, including voltage-gated K^+^ and Na^+^ channels.

To this end, we model a single channel as a two-state system that transitions between closed and open states. This reduced description is not intended to resolve the full molecular gating pathway; rather, it preserves the voltage-dependent occupancy of closed and open states. The channel gating is described by a reaction coordinate *ξ*, which summarizes the molecular configurations that distinguish closed and open channel states. The dynamics of *ξ* is governed by a voltage-dependent double-well potential, *U* (*ξ*) = *U* ^‡^(*ξ*^4^ − 2*ξ*^2^) − *Q*(*V* − *V*_G_)*ξ*; see Fig. 1c. Here, *U* ^‡^ is the barrier between the closed and open states, *Q* is the effective gating charge, and *V*_G_ is the gating potential at which the two states are equally favored. Crucially, *V* is the *local* trans-membrane voltage experienced by the channel; in general, *V* differs from *V*_ext_ because channel currents perturb the surrounding electrochemical environment. The local voltage *V* therefore tilts the potential and selects the thermodynamically favored channel state. The channel dynamics is modeled by the overdamped Langevin equation [11]

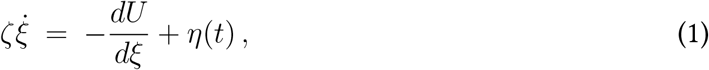

where *ζ* is the friction coefficient associated with the reaction coordinate and *η*(*t*) is thermal noise satisfying the fluctuation–dissipation relation ⟨*η*(*t*)*η*(*t*^′^)⟩ = 2*k*_B_*Tζ δ*(*t* − *t*^′^), with *T* being the temperature and *k*_B_ the Boltzmann constant. Equation (1) describes noisy motion on the gating landscape: the deterministic force relaxes the channel toward a local state, while thermal fluctuations occasionally drive barrier crossing.

Based on the conserved structure of the selectivity filter and the ion-selective transport properties of KcsA and KvAP channels [9, 10, 12], the current–voltage response of an open channel can be shown to follow a Goldman–Hodgkin–Katz-like law [13, 14] (see Supplemental Material (SM) Sec. I.2):

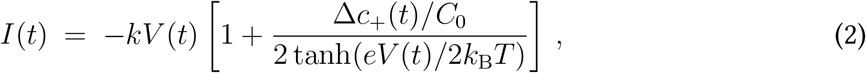

where *V* (*t*) and Δ*c*_+_(*t*) are, respectively, the local transmembrane voltage and potassium concentration difference across the membrane evaluated locally at the channel, *e* is the elementary charge, and *C*_0_ is the reference electrolyte concentration (∼ 150 mM). The parameter *k* is the intrinsic channel conductance.

Upon channel opening, the selective transport of cations breaks local electroneutrality in the electrolyte compartments and perturbs both the transmembrane potential and the local concentration difference. These perturbations arise from the formation of diffuse charge double layers of thickness of order the Debye length *λ*_D_ on either side of the membrane. For a constant source of current, the corresponding local voltage and concentration changes at the channel are 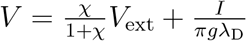∗, and 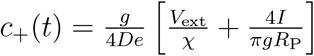, where *g* is the bulk electrolyte conductivity (*g* ≈ 1 S*/*m at physiological conditions), *D* is the potassium diffusivity, *χ* ≫ 1 is the ratio of the electrical double-layer capacitance to the membrane capacitance, and *R*_P_ is the effective pore radius [16]. Thus, even for a single channel, the voltage regulating the gating and current is not simply the externally imposed voltage, but includes the channel’s own electrochemical self-response. For external voltage differences comparable to the thermal voltage 2*k*_B_*T/e*, these relations in conjunction with Eq. (2) result in the following approximate expressions for the steady-state open probability and the current–voltage response as functions of the externally controlled voltage *V*_ext_ (SM Sec. I.3):

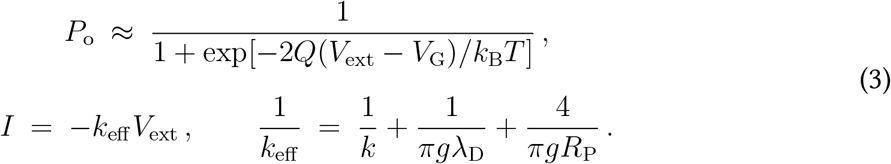

The effective single-channel conductance *k*_eff_ is smaller than the intrinsic conductance *k* because the current flow encounters additional resistances from the diffuse charge double layers and the finite pore geometry. These relations, together with fully nonlinear single-channel Langevin simulations, reproduce the measured KvAP open-probability and current–voltage curves reasonably well, as shown in Figs. 1d, e. This comparison calibrates the single-channel parameters used throughout this work, namely *Q* = 1.6*e, V*_G_ = −50 mV, *R*_P_ = 0.7 nm, *k* = 350 pS and an effective conductance *k*_eff_ ≈ 210 pS.

### Cooperativity of multiple channels

The selective cation current through an open channel not only modifies its own local transmembrane potential and concentration difference, but also perturbs the electrical environment experienced by other channels. This provides a physical mechanism for channel–channel coupling without requiring direct molecular contact between channels. For a membrane in an unbounded electrolyte, a localized channel current has recently been shown to generate spatiotemporal transmembrane-potential perturbations *V* (**x**, *t*) along the membrane through the charging and discharging of diffuse double layers on either side of the membrane [16]. After a short charging transient, over the distances most relevant for channel–channel coupling, this voltage perturbation has a long-ranged monopolar form, decaying as *V* ∼ 1*/r* with distance *r* from the open channel. Thus, the current through one open channel can alter the voltage experienced by nearby channels.

In the presence of electrodes or finite boundaries, this unbounded-electrolyte response is modified by confinement [17]. In particular, the long-ranged perturbation is cut off beyond the confinement length scale *L*, and the transmembrane potential is effectively screened at larger distances (Fig. 1f). Specifically, the voltage perturbation at any channel *j* due to a constant open current *I*_*i*_ through channel *i* is given by:

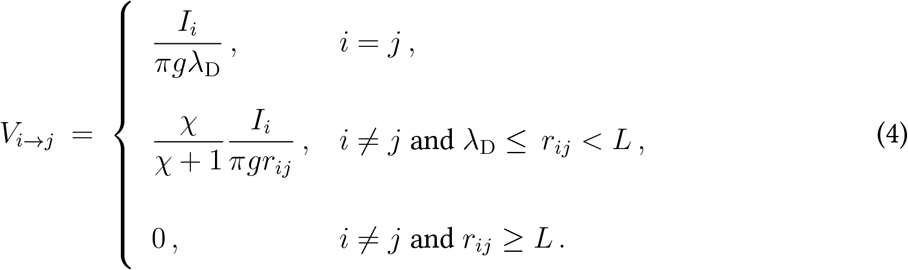

Here, *r*_*ij*_ is the distance between channels *i* and *j*. This expression captures the self-induced voltage shift of a channel through the local double-layer resistance, as well as the long-ranged monopolar perturbation affecting other channels up to the confinement scale *L*. The local voltage experienced by channel *j* is therefore 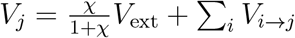, which is the voltage regulating the gating dynamics of channel *j*.

These voltage perturbations act as effective interaction terms between channels. When the gating voltage satisfies *V*_G_ < 0, the current produced by an open channel can make nearby channels experience a less negative local transmembrane potential than the externally imposed voltage alone. As a result, neighboring channels can be biased toward opening even when the applied voltage is below the isolated-channel activation voltage. For a membrane containing many channels, this feedback can therefore amplify channel opening and generate cooperative gating.

To understand the emergence of collective effects, we simulate the fully coupled nonlinear Langevin equations for a square lattice of *N* ion channels separated by a lattice spacing *ℓ* corresponding to a density *ρ* = 1*/ℓ*^2^, as shown schematically in Fig. 1a. These simulations are performed under periodic boundary conditions, wherein the system sizes, i.e., the numbers of ion channels, are chosen such that *Nℓ*^2^ > 4*L*^2^ to avoid interactions across periodic replicas. See SM Secs. II & III for a detailed description of the multichannel simulations including details on the channel-channel interactions and implementations using OpenMM [18].

Figure 1g shows the activation curves, measured by the mean open fraction *m*, at two different temperatures. Here, *m* = 0 corresponds to a state in which nearly all channels are closed, whereas *m* = 1 corresponds to a state in which nearly all channels are open. While isolated-channel gating occurs near the single-channel gating voltage *V*_ext_ = *V*_G_ ≃ −50 mV, the multichannel system exhibits collective activation at a substantially lower applied voltage. For example, for the simulations shown in Fig. 1g, where *L* = 27 nm and *ℓ* = 3 nm ^∗^, the collective activation occurs near *V*_ext_ ≃ −78 mV. Thus, the ensemble opens at voltages for which isolated channels would remain mostly closed. Moreover, the activation curves remain sigmoidal and sharpen as the temperature is lowered, indicative of an underlying phase transition between predominantly closed and predominantly open channel states, analogous to the sharpening of magnetization curves in interacting spin systems as a function of the applied magnetic field [19] or the volume of a liquid undergoing an equilibrium vapor-liquid transition as a function of pressure [20].

### Bistability and first-order phase transitions

In equilibrium systems, first-order phase transitions are characterized by discontinuous changes in an order parameter as a conjugate thermodynamic field is varied. A familiar example is the discontinuous change in density or volume across a liquid–vapor transition. While a true discontinuity exists only in the thermodynamic limit [19], finite systems still retain characteristic signatures of the transition, particularly in the scaling of the crossover behavior as the system passes from one phase to the other [21, 22]. As the system size increases, the crossover sharpens, and its width decreases toward zero in the thermodynamic limit.

For the multichannel system, we take the open fraction *m* as the order parameter distinguishing the collectively closed and collectively open states, and the external voltage *V*_ext_ as the conjugate field. Figure 2a shows the mean open fraction *m* as a function of *V*_ext_ for systems containing different numbers of ion channels. The system crosses over from a predominantly closed state to a predominantly open state as the external voltage is increased. This finite-size crossover is centered near the coexistence voltage *V* ^∗^(*T* ), where the collectively closed and open states are equally favored. The coexistence voltage *V* ^∗^(*T* ) ≃ −73.97 mV at *T* = 270 K is substantially more negative than the isolated-channel gating voltage *V*_G_ ≃ −50 mV, indicating the collective shift produced by current-induced electrical feedback. Moreover, increasing the number of channels sharpens the crossover around *V* ^∗^(*T* ), consistent with the approach to a discontinuity in the infinite-system limit.

**Figure 2:**
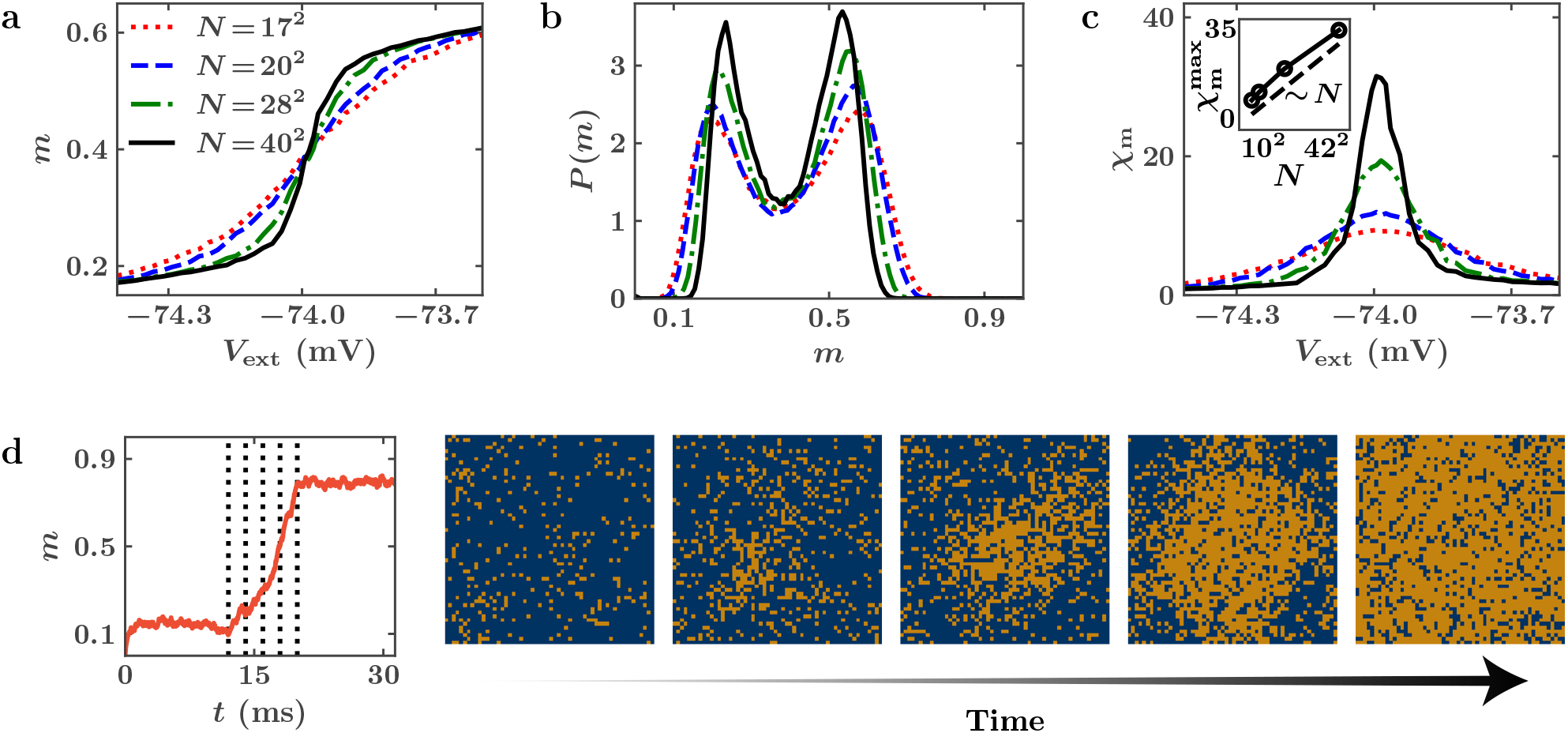
Evidence for first-order transitions. (a) The open-fraction response curve *m*(*V*_ext_) at *T* = 270 K sharpens with increasing system size around the activation threshold, corresponding to coexistence voltage *V* ^∗^(*T* ). (b) Near coexistence, *V*_ext_ ≃ − 73.97 mV, the order-parameter distribution is bimodal, with peaks corresponding to the collectively closed and collectively open states. The peak-to-minimum contrast increases with system size. (c) The susceptibility *χ*_*m*_ exhibits an increasingly sharp peak near the activation threshold. The peak value grows linearly with *N*, as shown in the inset, consistent with first-order transition scaling. (d) The multichannel system exhibits nucleation and growth dynamics characteristic of first-order transitions. A system initially maintained in a predominantly closed state at *V*_ext_ = −100 mV is stepped to *V*_ext_ = −72.36 mV at 230 K. The instantaneous open fraction *m*(*t*) remains near the metastable closed state before spontaneously growing toward the collectively open state. Snapshots at the five marked time points during the transition show that a predominantly open state nucleates and grows outward. Gold indicates open channels and blue indicates closed channels.

At coexistence, the two distinct phases coexist with their respective order-parameter values. The order-parameter distribution therefore exhibits bimodal behavior, with the two peaks corresponding to the two coexisting phases [21, 22]. In the present system, these peaks represent ensembles that are mostly closed or mostly open. These peaks are separated by a minimum at intermediate values of *m*, corresponding to configurations in which the two phases coexist and are separated by an interface. In an equilibrium first-order transition, the logarithmic difference between the peaks and the minimum grows as *N* ^(*d*−1)*/d*^, where *d* is the spatial dimension, owing to the finite thermodynamic cost of forming an interface between the two coexisting phases [19]. Figure 2b shows the distribution *P* (*m*) for the multichannel system at *T* = 270 K and for several system sizes, evaluated at their respective coexistence voltages *V* ^∗^(*T* ). The two peaks correspond to the coexisting phases, with mean open fractions *m*_1_ and *m*_2_. Increasing or decreasing the applied voltage away from *V* ^∗^(*T* ) by a small amount tilts the distribution toward the open or closed state, respectively, as shown in Fig. S6b. The peak-to-minimum difference increases with system size, consistent with an effective interfacial cost separating the collectively closed and open states.

A second finite-size signature is the enhancement of fluctuations near coexistence. The presence of two macroscopic states at coexistence, with respective mean numbers of open channels *M*_1_ = *Nm*_1_ and *M*_2_ = *Nm*_2_, implies that the mean-square fluctuations in the number of open channels scale as *N* ^2^ with system size. Consequently, the susceptibility *χ*_*m*_, which is the response of the open fraction to changes in the applied voltage, exhibits a peak at the coexistence voltage *V* ^∗^(*T* ) that must scale as *N* [19]. Figure 2c shows the susceptibility of the multichannel system, measured by the mean-square fluctuations *χ*_*m*_ := *N* ⟨[*m* − ⟨*m*⟩]^2^⟩, as a function of the external voltage. The susceptibility exhibits a peak near *V* ^∗^(*T* ), and the peak height 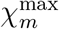 increases linearly with system size, indicating large collective fluctuations as the system switches from one state to the other, as expected for a first-order transition.

Lastly, first-order transitions are accompanied by metastability near coexistence. Upon a sudden change in the conjugate field, the system can persist for a finite time in the metastable phase until a critical cluster of the stable phase forms and grows [23, 24]. Indeed, the multichannel system exhibits such nucleation and growth behavior after a voltage quench from *V*_ext_ = −100 mV, where the channels are predominantly closed, to *V*_ext_ = −72.36 mV, just above the coexistence voltage at *T* = 230 K, as seen in Fig. 2d. Figure 2d shows a sequence of time snapshots of the multichannel dynamics, where a localized cluster of open channels (gold) nucleates within the closed background (blue) and then grows, eventually driving the system to the collectively open state. Taken together, the sharpening of the activation curve, the bimodal order-parameter distribution, the system-size scaling of the susceptibility peak, and the nucleation-and-growth dynamics provide evidence that the nonequilibrium multichannel system undergoes a bona fide first-order transition.

### Evidence for critical fluctuations and the critical point

In equilibrium systems, first-order transition lines often terminate at special points in the phase diagram, such as a triple point or a critical point. At a critical point, the collectively closed and collectively open phases become indistinguishable. As one approaches a critical point along a first-order transition line, the bimodal nature of the order-parameter distribution disappears, reflecting the vanishing interfacial cost between the two coexisting phases. In the vicinity of the critical point, the system is therefore dominated by large fluctuations, where small changes in the applied field result in strong responses [19].

Figure 3a shows a snapshot of a large multichannel system with *N* = 6400 channels at *T* = 280 K and *V*_ext_ = −74.19 mV, near the putative critical point. The spatial distribution of closed and open channels shows large fluctuations, with extended regions of both states appearing throughout the system. To identify the critical point of the multichannel system, we simulate the system at several temperatures and locate the coexistence voltage *V* ^∗^(*T* ); see SM Fig. S6c and SM Sec. IV for details. We do this by identifying the external voltage at which the two peaks in the order-parameter distribution *P* (*m*) have equal height. Figure 3b shows *P* (*m*) at *V* ^∗^(*T* ) for increasing temperature. As the temperature is increased, the distribution loses its bimodal structure and evolves from two peaks into a single broad peak, indicating the disappearance of the distinction between the collectively closed and collectively open states. This behavior suggests the existence of a critical point near *T* ≃ 283 K.

**Figure 3:**
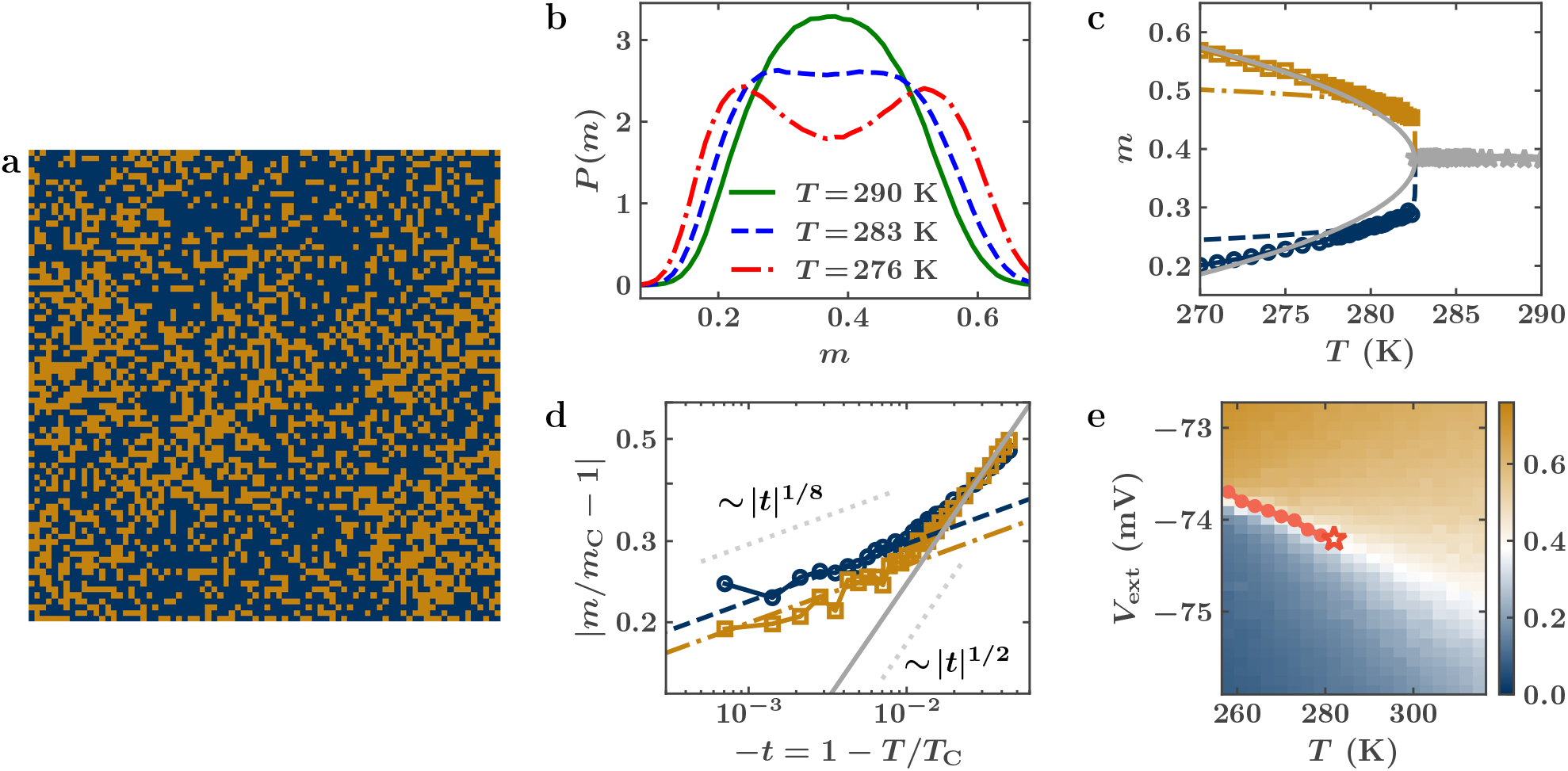
Near-critical behavior and the critical point. (a) Snapshot of an 80 × 80 multichannel system at *T* = 280 K and *V*_ext_ = −74.19 mV, near the identified critical point. Large patches of collectively open or collectively closed channels appear throughout the system, consistent with an increasing correlation length near criticality. Gold indicates open channels and blue indicates closed channels. (b) Order-parameter distributions *P* (*m*) at the coexistence voltage *V* ^∗^(*T* ) for three temperatures: below *T*_C_, where the distribution is bimodal; near *T*_C_, where the distribution broadens; and above *T*_C_, where the distribution exhibits a single peak. (c) Mean steady-state open fraction *m* as a function of temperature. For *T* < *T*_C_, the two branches correspond to the mostly closed state (blue) and mostly open state (gold), identified from the two peaks of *P* (*m*) at coexistence. For *T* > *T*_C_, the single branch corresponds to the single peak of *P* (*m*). The dashed blue and gold curves show fits to (*T*_C_ − *T* )^1*/*8^, while the solid gray curve shows the expected meanfield scaling (*T*_C_ − *T* )^1*/*2^, which deviates from the simulations near the critical point. (d) Data from panel (c) plotted in terms of the reduced temperature −*t* = 1 −*T/T*_C_ and the reduced open fraction |*m/m*_C_ −1| on logarithmic axes, showing consistency with the 1*/*8 order-parameter exponent of the two-dimensional Ising universality class. A crossover toward mean-field behavior occurs for −*t* ≳ 0.01. (e) Numerical phase diagram in the applied-voltage–temperature plane. Gold and blue indicate predominantly open and predominantly closed states, respectively. The coexistence curve is shown in red and terminates at the identified critical point, marked by a star, with *V*_C_ = −74.23 mV and *T*_C_ = 282.6 K. All critical values are functions of the chosen single-channel parameters.

Another important characteristic of the critical point is the variation of the order parameter in its vicinity. Figure 3c shows the mean open fractions in the two coexisting phases, corresponding to the two peaks of the bimodal distribution *P* (*m*) at the coexistence voltage *V* ^∗^(*T* ), as a function of temperature. At higher temperatures, where the distribution has a single peak for all *V*_ext_, we plot the single mean open fraction at the voltage where the susceptibility is at its maximum. As *T* approaches *T*_C_, the two branches approach one another, indicating that the collectively closed and collectively open phases become indistinguishable at the critical point. Near criticality, this approach is expected to follow a power law |*m*(*T* ) − *m*_C_| ∼ |*T*_C_ − *T* |^*β*^ . As shown in Fig. 3d, the rescaled order parameter is fit well by *β* = 1*/*8, the order-parameter exponent of the equilibrium two-dimensional Ising universality class [25]. The appearance of the two-dimensional Ising exponent is consistent with universality: near a critical point, scaling behavior is determined by system-wide features such as dimensionality, symmetry of the order parameter, and range of interactions, rather than by microscopic details. Thus, although ionic currents hold the ensemble in a nonequilibrium steady state, the observed scaling suggests that the nonequilibrium drive may be irrelevant under coarse-graining for this non-conserved scalar order parameter, leaving the transition in the equilibrium Ising universality class [26]. A rigorous demonstration of this irrelevance, for instance via an explicit renormalization-group analysis of the coarse-grained dynamics, is left to future work. Nevertheless, this analysis yields a critical temperature *T*_C_ = 282.6 K.

While this work establishes the order-parameter scaling near the critical point, other critical signatures may also be examined, including the divergence of the susceptibility and the slowing down of relaxation dynamics as *T*_C_ is approached. Analyzing these divergences requires considerably larger simulations and longer sampling times, and we leave this endeavor for future work. Nevertheless, the observed order-parameter scaling provides evidence that the collective behavior of the multichannel system follows Ising-like criticality near the critical point. Figure 3e summarizes the resulting voltage–temperature phase diagram, showing a first-order transition line separating the collectively closed and collectively open states and terminating at a critical point.

### A nonequilibrium dimensionless parameter controls the critical temperature

The critical temperature in the *V*_ext_–*T* phase diagram indicates the temperature scale around and below which channel ensembles activate cooperatively. When the physiological temperature is much greater than the collective critical temperature *T*_C_, thermal fluctuations overwhelm the voltage-induced coupling between channels and each channel effectively gates independently. Therefore, the value of *T*_C_ determines whether collective channel activation is expected to be relevant under physiological conditions. This raises the question as to what controls *T*_C_.

Given the existence of phase-transition behavior and its similarity to equilibrium phase transitions in Figs. 2 and 3, the fully coupled Langevin equations are amenable to a mean-field analysis. For applied voltages on the order of the thermal voltage, this analysis leads to the following expressions for the critical temperature and critical applied voltage (SM Sec. V):

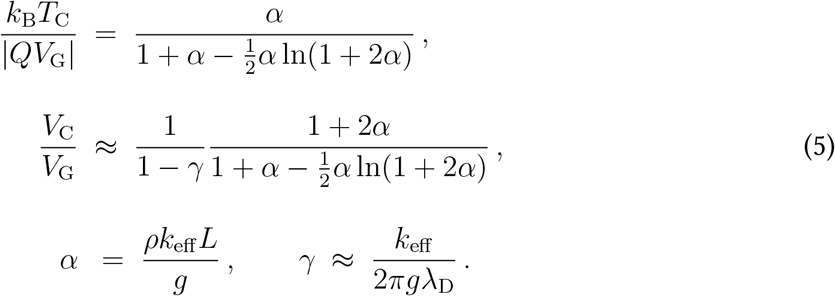

The parameters *α* and *γ* are dimensionless conductance ratios that compare ionic transport through the membrane with transport through the surrounding electrolyte, analogous to Biot numbers in heat transfer [27]. *ρ* = 1*/ℓ*^2^ is the areal channel density, where *ℓ* is the inter-channel spacing. The parameter *α* is the primary control parameter for collective behavior. Specifically, *α* can be written as *α* ≈ (*ρk*_eff_ )*/*(*g/L*), and therefore measures the ratio of the effective conductance of ions per unit area through the membrane, set by the channel conductance and channel density, to the conductance per unit area through the surrounding electrolyte over the confinement length *L*. The key quantity is therefore not channel density alone, but the conductance of the membrane relative to the ability of the surrounding electrolyte to redistribute the resulting current. Increasing the membrane conductance or decreasing the channel spacing increases the current density through the membrane, leading to stronger voltage perturbations and larger cooperative effects. Conversely, increasing the bulk electrolyte conductivity or decreasing the confinement length reduces the voltage perturbations in the surrounding electrolyte and therefore weakens cooperative gating.

The second parameter, *γ*, is the ratio of the effective channel conductance to the access-like conductance of the electrolyte near the channel pore. It quantifies how efficiently ions are transported through the pore relative to how they are supplied to and removed from the pore by the surrounding electrolyte. Under physiological conditions, *γ* ≪ 1, indicating that conductance through the pore is typically the rate-limiting step. Thus, the dimensionless number *α* primarily governs the importance of collective behavior in biological systems: when *α* ≪ 1, channels gate nearly independently, whereas when *α* ≈ 1, current-induced voltage perturbations become strong enough to produce cooperative gating and a finite critical temperature comparable to physiological temperature.

### Collective channel activation in biological excitable membranes

The role of voltage-dependent, ion-selective conductances in electrical signaling was established more than seventy years ago by Hodgkin and Huxley through their theory of the action potential, which identified distinct dynamic roles for Na^+^ and K^+^ currents in the squid giant axon [1]. This ion-selective permeability is now understood to arise from voltage-gated Na^+^ and K^+^ channels. Given the bio-electrical phase transitions identified here, it is natural to ask whether the emergent cooperativity described in this work may be relevant to the collective behavior of Na^+^ and K^+^ channels underlying action potentials in the squid giant axon. We use this system as a first order-of-magnitude test because many of the relevant electrophysiological parameters have been measured directly.

For the squid giant axon, the relevant confinement length is set by the finite axonal cross-section rather than by bounding electrodes. In a cylindrical membrane, the response to a localized current crosses over from a monopolar perturbation to a screened field over axial distances of order the axon diameter [28]; we therefore take *L* ≈ 550 *µ*m. The Na^+^ channel density has been estimated to lie in the range 166–553 *µ*m^−2^ [29, 30]. Taking the lower estimate *ρ*_Na_ ≈ 166 *µ*m^−2^, together with an effective single-channel conductance *k*_eff_ ≈ 14 pS [31], we estimate the conductance ratios *α* and *γ* under physiological conditions. For axons treated by pronase, which enzymatically abolishes fast Na^+^-channel inactivation and thereby isolates activation gating, the Na^+^ channel gating charge and gating potential have been estimated as *Q* ≈ 2*e* and *V*_G_ ≈ −20 mV [32]; see also single channel measurements in [33]. The resulting *T*_C_ is controlled by the bulk conductivity through *α* ∝ 1*/g*. In the classical measurements by Hodgkin and Huxley [1], the axoplasm has an estimated conductivity of *g* ≈ 2.8 S*/*m and the external sea-water is comparable, *g* ≈ 4 S*/*m. Taking *g* ≈ 3 S*/*m and *λ*_D_ ≈ 0.8 nm gives *α* ≈ 0.43–1.42 and *γ* ≈ 0.00093 across the cited density range, and a Na^+^ channel critical temperature *T*_C_ ≈ 150– 450 K, together with a critical voltage *V*_C_ ≈ −30 to −54 mV. The crossover density at which *T*_C_ reaches the squid physiological temperature (∼ 283 K) is *ρ*^∗^ ≈ 340 *µ*m^−2^, which lies within the measured density range. Thus, one of the most thoroughly characterized excitable membranes appears to lie near the threshold at which the collective mechanism described here becomes physiologically relevant. Accurate resolution of the collective activation threshold therefore requires further careful measurements of the Na^+^ channel density in the squid giant axon.

The corresponding estimate for K^+^ channels yields a different result. Experiments indicate an effective single-channel conductance *k*_eff_ ≈ 17.4 pS [34] and a channel density *ρ*_K_ ≈ 67 *µ*m^−2^ [35], substantially lower than that of Na^+^ channels. Because explicit measurements of the relevant K^+^ channel gating charge and gating potential in this system are less direct, we estimate the critical temperature assuming values comparable to those of Na^+^ channels, namely *Q* ≈ 1.6*e*–2*e* and *V*_G_ ≈ −20 mV. These estimates, at the same *g* ≈ 3 S*/*m, yield *α* ≈ 0.21 and 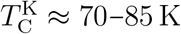, well below physiological temperature. This comparison suggests that, while Na^+^ channels in the squid giant axon straddle the collective threshold, K^+^ channels are expected to gate more independently, primarily because of their lower surface density and correspondingly smaller value of *α*.

Another system of interest is the axon initial segment (AIS), a specialized neuronal region known to initiate action potentials and enriched in voltage-gated Na^+^ channels [4]. Areal Na^+^ channel conductances have been experimentally estimated to be 2500 pS*/µ*m^2^ in cortical neurons [4], and are sometimes taken to be as large as 30000 pS*/µ*m^2^ in models describing their firing patterns [36]. Such high channel densities indicate the possibility that collective activation may occur in the AIS. Indeed, cooperative Na^+^ channel activation has been invoked as a possible mechanism for the rapid onset of cortical spikes [37], although alternative explanations based on the neuronal spatial geometry, spike initiation properties, and AIS compartmentalization and spatial organization have also been proposed [38–40]. Using the present theory, with Nav1.6 gating parameters *Q* ≈ 2.5*e, V*_G_ ≈ −36 mV [41–43], a mammalian conductivity *g* ≈ 1 S*/*m, an upper estimate of the areal Na^+^ channel conductance *ρk*_eff_ ≈ 30000 pS*/µ*m^2^, and a relatively large axonal diameter *L* ≈ 3 *µ*m, we obtain *α* ≈ 0.09 and a Na^+^ channel critical temperature *T*_C_ ≈ 87 K, well below physiological temperature. Using lower experimental areal conductance values would only reduce this estimate. Thus, despite the high Na^+^ channel density in the AIS, the much smaller confinement length strongly suppresses the current-induced electrical feedback, suggesting that spike initiation in the AIS is unlikely to arise from the collective phase-transition mechanism described here. This conclusion does not preclude other forms of channel cooperativity contributing to rapid spike onset.

The nodes of Ranvier in mammalian myelinated nerve fibers provide another example of a Membrane region with very high Na^+^ channel density. Early toxin-binding studies of rabbit sciatic nerve estimated Na^+^ channel numbers of order 7.2 × 10^5^ channels per node, corresponding to a density of approximately 12000 *µ*m^−2^ for a nodal area of about 60 *µ*m^2^ [5]. Although later electrophysiological studies provided lower estimates, roughly 1500–2000 *µ*m^−2^ [44], nodes of Ranvier remain among the highest-density Na^+^ channel assemblies and are therefore natural candidates for collective activation. However, as illustrated by the axon initial segment, high channel density alone is not sufficient: a small axonal diameter lowers the conductance ratio *α* and weakens the cooperative effect. Nodes of Ranvier in large-diameter peripheral nerve fibers therefore present a particularly intriguing possibility. For rabbit sciatic nerve, using the reported nodal area and a nodal length of 1–2 *µ*m gives an effective fiber diameter of ≈ 10.5–21 *µ*m [5]. Taking a conservative Na^+^ channel density *ρ*_Na_ ≈ 2500 *µ*m^−2^, an effective conductance *k*_eff_ ≈ 14 pS, Nav1.6 gating parameters *Q* ≈ 2.5*e*, and *V*_G_ ≈ −36 mV [41–43], and a mammalian conductivity *g* ≈ 1 S*/*m we obtain *T*_C_ ≈ 303–547 K as the nodal length is varied over 1–2 *µ*m (the larger diameter, for the shorter 1 *µ*m node, giving the higher *T*_C_). This range straddles the mammalian physiological temperature (∼ 310 K), lying just below it for a 2 *µ*m node and well above it for a more typical 1 *µ*m node. Correspondingly, the crossover density shifts from *ρ*^∗^ ≈ 2800 *µ*m^−2^ at 2 *µ*m to ≈ 1400 *µ*m^−2^ at 1 *µ*m, bracketing the conservative density estimate of 2500 *µ*m^−2^. Among the systems considered here, nodes of large-diameter peripheral fibers are therefore the most promising candidates for collective Na^+^ channel activation under physiological conditions.

While the cases discussed above suggest that collective activation may occur in realistic biological systems, it is desirable to obtain a direct experimental observation of the phase transition described here. Reconstituted membrane systems of the kind used in Ref. [9] may provide a natural experimental testbed, since they allow single-channel electrophysiological measurements and could be further engineered to vary the density of incorporated channels. A prominent signature of emergent cooperativity, and possibly of an underlying phase transition, would be a systematic shift in the activation threshold of a multichannel system relative to the corresponding single-channel activation curve, as illustrated in Fig. 1g. Such an observation would not only test the possibility of bioelectrical phase transitions in ion-channel ensembles, but also directly probe the electrical feedback mechanism arising from the long-ranged monopolar transmembrane-potential perturbations generated by ion-selective channel currents, as described by Eq. (4).

## Acknowledgments

J.B.F. acknowledges support from the U.S. Department of Energy, Office of Science, Office of Advanced Scientific Computing Research, through the Department of Energy Computational Science Graduate Fellowship under Award No. DE-SC0023112. K.K.M. and J.B.F. acknowledge support from the Director, Office of Science, Office of Basic Energy Sciences, of the U.S. Department of Energy under Contract No. DE-AC02-05CH11231. K.S. acknowledges support from the McKnight Foundation, Sloan Foundation, Philomathia Foundation, and the University of California, Berkeley. K.S. and H.R. acknowledge support from the National Institutes of Health (U01NS136405). This research used resources of the National Energy Research Scientific Computing Center, a U.S. Department of Energy Office of Science User Facility located at Lawrence Berkeley National Laboratory, under NERSC Award No. BES-ERCAP0023682. We thank Prof. David Limmer for useful discussions.

## Contribution Statement

K.K.M. and K.S. conceived the project. All authors designed and performed the research. J.B.F. wrote the software and performed the simulations. All authors contributed to the analysis of the data, interpretation of the results, and writing of the paper. K.S. and K.K.M. supervised the project.

∗ Given *χ* ≫ 1, almost all of the applied potential *V*_ext_ is borne by the membrane [15].

∗ These are chosen to make direct simulations of the long-ranged interacting system computationally tractable when establishing the existence of the phase transition; the mean-field theory we develop below extends the results to physiological values of *L* and *ℓ*.

