## Supplemental Material for "Bioelectrical phase transitions"

#### Non-equilibrium bioelectrical phase transitions

**References** **33**

This Supplemental Material is organized as follows. In Sec. I, we describe the model of a single ion channel and discuss the results of single-channel simulations. In Sec. II, we model the interactions between ion channels and develop a simplified model for a system of multiple channels. In Sec. III, we present the numerical methods used to simulate the stochastic dynamics of a system of multiple channels, and in Sec. IV, we describe how we analyze the simulation data to construct the applied voltage–temperature phase diagrams. Finally, in Sec. V, we develop a mean-field theory to describe the collective behavior of many channels and provide the phase diagrams, along with analytical expressions for the critical voltage and temperature in the mean-field limit.

### I. Single channel behavior

We describe the behavior of a single ion channel by accounting for the stochastic gating dynamics, the transmembrane ionic current through the channel, the long-ranged electrochemical perturbation generated by the current, and the combined effect of these processes.

#### 1. Open-to-closed transition

We consider a reduced channel-gating model with two states: open and closed. In constructing the dynamics of the single channel, we neglect inactivation, which is known to occur in voltage-gated ion channels [1–3]; that is, the ability of the channel to transition from an open state to an inactive state that is unable to transmit transmembrane current. At present, it is our desire to study the nature of collective activation in ion-channel ensembles. Incorporating inactivation presents an avenue for further study.

For simplicity, we model the transition between the closed and open states as a single barrier-crossing process. We introduce a one-dimensional reaction coordinate $\xi$ and assign the closed and open states to $\xi = -1$ and $\xi = +1$, respectively. This bistability is represented by a symmetric double-well free energy,

$$U_0(\xi) \ = \ U^{\ddagger}\left(\xi^4 - 2\xi^2\right) \ , \tag{S.1}$$

where $U^{\ddagger}$ sets the barrier height at $\xi = 0$. The local minima of the free energy are located at $\xi = -1$ and $\xi = +1$. Although $\xi$ is defined over $(-\infty, \infty)$, the system typically remains localized near one well, with the states classified as closed for $\xi < 0$ and open for $\xi \geq 0$.

The symmetric double-well potential in Eq. (S.1) serves as an unbiased reference in which the closed and open states are equiprobable. For a voltage-gated channel, this equiprobable condition occurs when the membrane potential $V$ equals the gating voltage $V_{\mathrm{G}}$. Away from this value, the membrane potential tilts the landscape such that $V > V_{\mathrm{G}}$ favors opening and $V < V_{\mathrm{G}}$ favors closing. We model this voltage-dependent asymmetry by adding a linear bias to the double-well free energy,

$$U(\xi) \ = \ U^{\ddagger}\left(\xi^4 - 2\xi^2\right) - Q\,(V - V_{\mathrm{G}})\,\xi \ , \tag{S.2}$$

where $Q$ is the voltage sensitivity of the gating coordinate, measured in units of charge. For $Q(V - V_{\mathrm{G}}) \ll U^{\ddagger}$, the minima do not move significantly from $\xi = \pm 1$, and the energy difference between the closed and open states is $\approx 2Q(V - V_{\mathrm{G}})$. We can associate $2QV$ with the energy change due to an applied field, where a conformational change in the protein moves charged residues within the applied field. The other term, $-2QV_{\mathrm{G}}$, corresponds to the baseline difference in energy due to structural differences. Higher-order terms may enter more detailed descriptions of gating, but the linear term suffices to capture the experimentally observed voltage dependence of the open probability, as we show later.

We model the evolution of the reaction coordinate in the free-energy landscape in Eq. (S.2) using the overdamped Langevin equation,

$$\zeta\dot{\xi} \ = \ -\frac{dU}{d\xi} + \eta(t) \ , \tag{S.3}$$

where $\zeta$ is the drag coefficient. The thermal noise $\eta(t)$ has zero mean, $\langle \eta(t) \rangle = 0$, and covariance $\langle \eta(t)\eta(t') \rangle = 2\zeta k_{\mathrm{B}} T\,\delta(t - t')$, as required by the fluctuation-dissipation theorem. Here, $\langle \cdot \rangle$ denotes an average over realizations of the thermal noise.

Within the physiological range of membrane potentials, the bias remains small compared with the barrier height, i.e., $|a| \ll 1$ with $a \equiv Q(V - V_G)/U^\ddagger$, and the closed and open states remain well separated by the energy barrier. As argued in the following Sec. I.1 (a), the barrier height is also large compared with thermal energy, i.e., $\beta U^\ddagger \gg 1$, where $\beta \equiv (k_B T)^{-1}$. Under these conditions, Kramers' theory [4] provides the transition timescales between the two states:

$$\tau_{c\to o} = \frac{\pi\zeta}{2\sqrt{2}U^\ddagger}\left(1 + \frac{3a}{16} + \mathcal{O}(a^2)\right) e^{\beta U^\ddagger(1-a+\mathcal{O}(a^2))} , \quad (S.4)$$

$$\tau_{o\to c} = \frac{\pi\zeta}{2\sqrt{2}U^\ddagger}\left(1 - \frac{3a}{16} + \mathcal{O}(a^2)\right) e^{\beta U^\ddagger(1+a+\mathcal{O}(a^2))} . \quad (S.5)$$

Here, $\tau_{c\to o}$ is the mean time for a transition from the closed to the open state by crossing the energy barrier $U^\ddagger$, and $\tau_{o\to c}$ is the corresponding mean time for the reverse transition. Thus, on average, a channel remains closed for $\tau_{c\to o}$ before opening and remains open for $\tau_{o\to c}$ before closing. Since $\beta U^\ddagger \gg 1$, even a small bias satisfying $|a| \ll 1$ can significantly affect the transition times through the exponential term. We therefore keep the $\mathcal{O}(a)$ correction in the exponent while neglecting the corresponding correction in the prefactor:

$$\tau_{c\to o} = \tau_G e^{-\beta Q(V-V_G)} , \qquad \tau_{o\to c} = \tau_G e^{\beta Q(V-V_G)} , \quad (S.6)$$

where

$$\tau_G \equiv \frac{\pi\zeta}{2\sqrt{2}U^\ddagger} e^{\beta U^\ddagger} , \quad (S.7)$$

is the single-channel mean transition timescale for the unbiased case $a = 0$.

For a fixed membrane potential $V$, the steady-state open probability is determined by the relative residence times in the open and closed states. Using the approximate transition timescales derived above, we obtain

$$P_o = \frac{\tau_{o\to c}}{\tau_{o\to c} + \tau_{c\to o}} = \frac{1}{1 + e^{-2\beta Q(V-V_G)}} . \quad (S.8)$$

#### (a). Estimation of drag coefficient and energy barrier

We provide an estimate of the drag coefficient $\zeta$ and the barrier $U^\ddagger$ governing the single channel dynamics in Eq. (S.3).

In voltage-gated potassium channels, gating is controlled by a voltage-sensing domain whose charged residues respond to the local electric field [5]. In our reduced two-state model, this *voltage sensor* occupies different depths in the membrane in the closed and open states, resulting in the electrostatic bias introduced above. The actual conformational change between the closed and open states involves a rotary-like motion of the protein, which gates the channel pore [6]. As a crude approximation, we describe this motion as rotational diffusion. For a cylindrical protein of radius $R$ in a membrane of thickness $\delta$, the Saffman-Delbrück rotational drag coefficient [7] is

$$\zeta = 4\pi\nu R^2 \delta , \quad (S.9)$$

where $\nu$ is the membrane (shear) viscosity. Taking $\delta = 4\,\text{nm}$, $R \approx 0.9\,\text{nm}$, and $\nu \approx 0.01\,\text{Pa}\cdot\text{s}$, we obtain $\zeta \approx 2.5\,\text{eV}\cdot\text{ns}$. This drag coefficient sets a timescale,

$$\tau_\xi = \beta\zeta \approx 100\,\text{ns} , \quad (S.10)$$

where we use room temperature $T = 298\,\text{K}$, so that $\beta^{-1} = 25\,\text{meV}$. This timescale represents the bare diffusive timescale of the gating variable and characterizes how quickly relaxation occurs toward the local equilibrium within an energy well. Comparing $\tau_\xi$ with the unbiased gating timescale $\tau_G$ yields

$$\frac{\tau_G}{\tau_\xi} = \frac{\pi}{2\sqrt{2}} \frac{e^{\beta U^\ddagger}}{\beta U^\ddagger} , \quad (S.11)$$

which indicates that the gating timescale is exponentially longer than the conformational relaxation timescale when $\beta U^\ddagger \gg 1$. This implies that barrier crossing is a rare event and that the gating variable relaxes locally in each well much more quickly than it moves between conformations.

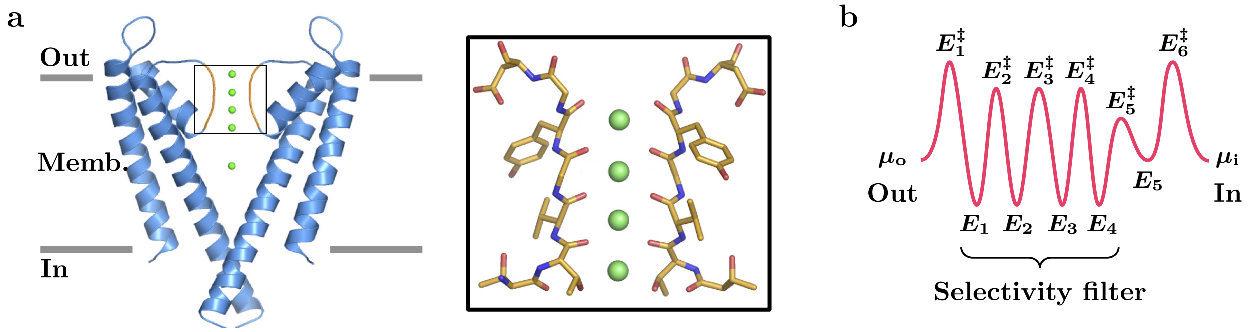

Figure S1: (a) The cryo-EM-determined structure of the KcsA channel, with the inset showing the selectivity filter (i.e., the pore) of the channel. This filter is such that the channel has high selectivity for potassium ions on the basis of charge and size. The green spheres represent potassium ions in the four binding sites of the channel. Figure adapted from Fig. 1 of Ref. [13]. (b) A schematic of the approximate energy landscape for the traversal of a potassium ion through the pore of a potassium channel, with position along the channel serving as the implied reaction coordinate. The selectivity filter is represented by four equal-depth wells, with the fifth well corresponding to the channel's cavity.

We next estimate the energy barrier between the closed and open states. Computational studies using molecular dynamics simulations have attempted to compute free-energy profiles for the gating transition and thereby infer effective barrier heights [8, 9]. Reported values vary widely, with $\beta U^\ddagger$ ranging from 50 to 120. From Eqs. (S.6) and (S.7), such barriers would imply transition times exceeding $10^6$ years, a discrepancy noted in these studies as well.

Accordingly, we estimate the effective barrier by fitting the gating timescale $\tau_G$ to experimental measurements and then using Eq. (S.11), together with our estimate of $\tau_\xi$, to infer the barrier height $\beta U^\ddagger$. This strategy has been used in experimental studies [10, 11]. The channel activation timescale $\tau_G$ at room temperature has been reported to range from $10\,\mu$s for rapidly activating channels [11] to 1 ms for slower channels [12]. Together with our estimate $\tau_\xi \approx 100$ ns, this range yields $\beta U^\ddagger \sim 6$–11. We use $U^\ddagger \approx 0.2$ eV, corresponding to $\beta U^\ddagger = 8$ at room temperature, which allows sufficiently fast equilibration while maintaining a robust energy barrier well above thermal energy. This choice gives an unbiased gating timescale of $\tau_G \approx 40\,\mu$s.

#### 2. Constitutive model for current

We next describe how the ionic current through an open channel depends on the electrochemical driving force across the membrane, which is determined by the membrane potential and the concentration difference. To construct this constitutive relation, we consider the channel pore structure that mediates ion transport. X-ray crystallography and cryo-electron microscopy have resolved potassium channel pore structures, as illustrated in Fig. S1a for the KcsA channel. Although the KcsA channel is not voltage-gated, its pore structure is conserved across the potassium channel family [5, 12, 14, 15]. Within the pore, the selectivity filter (Fig. S1a inset) contains binding sites for potassium ions, which reduce the energetic cost of desolvation from bulk solution. Motivated by this structure, we describe ion motion through the pore using an effective energy landscape with multiple barriers, as schematized in Fig. S1b. Here, we treat the pore structure, and therefore the energy landscape, as independent of ion occupancy.

We further simplify the model by considering the low-occupancy limit of the pore. In this limit, multiple ions are rarely present simultaneously, allowing us to neglect ion–ion interactions such as electrostatic repulsion and steric exclusion between ions. Although experimentally resolved occupancy patterns [16] and computational studies in physiological solutions [17] suggest that potassium channels can contain multiple ions under physiological conditions, this simplified description provides a useful starting point for constructing a constitutive model and, as shown below, leads to a familiar expression for the channel current.

We model ion transport inside the pore as transitions among the $N-1$ internal sites, with $x_i$ denoting

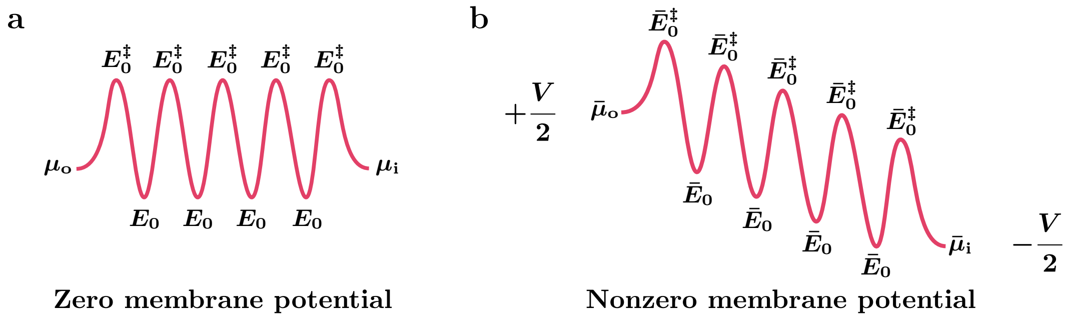

Figure S2: (a) An approximate energy landscape in which all well depths are equal and equally spaced along the reaction coordinate, with barriers of equal height. (b) Applying a membrane potential tilts the entire energy landscape in the direction of lower potential due to the charge of the potassium ion.

the average occupancy of site $i$ for $i = 1, \ldots, N-1$. Each transition is described by first-order kinetics: the transfer rate from site $i$ to $i \pm 1$ is $k_{i\to i\pm1} x_i$, where $k_{i\to i\pm1}$ is the associated rate constant. The ion-transport dynamics are then described by the master equation [18]:

$$\frac{d\mathbf{x}}{dt} = -\mathbf{W}\mathbf{x} + \mathbf{r} , \tag{S.12}$$

where $\mathbf{x} = [x_1, x_2, \ldots, x_{N-1}]^T$ is the vector of site occupancies and $\mathbf{r} = [r_{0\to1}, 0, \ldots, 0, r_{N\to N-1}]^T$ contains the boundary terms. Here, $r_{0\to1}$ and $r_{N\to N-1}$ represent entrance rates from the two boundaries into sites 1 and $N-1$, respectively. The rate matrix $\mathbf{W}$ is defined by

$$\mathbf{W} = \begin{bmatrix} k_{1\to0} + k_{1\to2} & -k_{2\to1} & & \\ -k_{1\to2} & k_{2\to1} + k_{2\to3} & -k_{3\to2} & \\ & & \ddots & \\ & & -k_{N-2\to N-1} & k_{N-1\to N-2} + k_{N-1\to N} \end{bmatrix} , \tag{S.13}$$

where the diagonal entries encode outgoing rates from each internal site, while the off-diagonal entries encode incoming rates from neighboring sites.

#### (a). Steady-state current

We first analyze the steady state, corresponding to $d\mathbf{x}/dt = \mathbf{0}$ in the master equation (S.12). In steady state, there is no accumulation at each site, and therefore the net forward flux $(i \to i+1)$ between each pair of neighboring sites must be equal. Solving the resulting linear system, we obtain this net forward flux in steady state as

$$j_{\mathrm{SS}} = \frac{\left(\prod_{i=1}^{N-1} \frac{k_{i\to i+1}}{k_{i\to i-1}}\right) r_{0\to1} - r_{N\to N-1}}{1 + \sum_{j=1}^{N-1} \left(\prod_{i=j}^{N-1} \frac{k_{i\to i+1}}{k_{i\to i-1}}\right)} , \tag{S.14}$$

written in terms of ratios between forward and backward rate constants. To evaluate these ratios, we first consider a neutral solute and determine the rate constants from the energy landscape inside the pore using Kramers' theory [4]. Electrostatic contributions are then included to obtain the analogous result for charged ions.

The transition rates out of site $i$ are given by

$$k_{i\to i-1} = \frac{\beta D \omega_i^\ddagger \omega_i}{2\pi} e^{-\beta(E_i^\ddagger - E_i)} , \qquad k_{i\to i+1} = \frac{\beta D \omega_{i+1}^\ddagger \omega_i}{2\pi} e^{-\beta(E_{i+1}^\ddagger - E_i)} , \tag{S.15}$$

where $D$ is the solute diffusivity, $E_i$ denotes the minimum of the energy well at site $i$, and $E_i^\ddagger$ denotes the barrier between sites $i-1$ and $i$. Here, $\omega_i$ and $\omega_i^\ddagger$ denote the frequency factors associated with the local curvatures of the well and the barrier, respectively. Taking the ratio of the two transition rates eliminates the dependence on the well properties, $E_i$ and $\omega_i$:

$$\frac{k_{i\to i+1}}{k_{i\to i-1}} = \frac{\omega_{i+1}^\ddagger}{\omega_i^\ddagger} e^{-\beta(E_{i+1}^\ddagger - E_i^\ddagger)} \ . \tag{S.16}$$

The product appearing in Eq. (S.14) then telescopes to

$$\prod_{i=j}^{N-1} \frac{k_{i\to i+1}}{k_{i\to i-1}} = \frac{\omega_N^\ddagger}{\omega_j^\ddagger} e^{-\beta(E_N^\ddagger - E_j^\ddagger)} \ . \tag{S.17}$$

It now remains to specify the boundary entrance rates $r_{0\to 1}$ and $r_{N\to N-1}$. We treat the two sides of the membrane as reservoirs with chemical potentials $\mu_\mathrm{o}$ and $\mu_\mathrm{i}$ for the transported solute, representing the outside and inside of the cell. Applying Kramers' theory at the two pore entrances yields

$$r_{0\to 1} = \frac{\beta D \omega_1^\ddagger \omega}{2\pi} e^{-\beta(E_1^\ddagger - \mu_\mathrm{o})} \ , \qquad r_{N\to N-1} = \frac{\beta D \omega_N^\ddagger \omega}{2\pi} e^{-\beta(E_N^\ddagger - \mu_\mathrm{i})} \ , \tag{S.18}$$

where $\omega$ is the boundary frequency factor, assumed to be the same on either side of the membrane. Substituting these expressions into Eq. (S.14) yields the steady-state flux,

$$j_\mathrm{SS} = \frac{\beta D \omega}{2\pi} \frac{e^{\beta\mu_\mathrm{o}} - e^{\beta\mu_\mathrm{i}}}{\sum_{j=1}^{N} \frac{1}{\omega_j^\ddagger} e^{\beta E_j^\ddagger}} \ . \tag{S.19}$$

This expression shows that the steady-state flux is controlled by the local maxima of the energy landscape, corresponding to the transition barriers along the pore, rather than by the depths of the wells. The flux is driven by the jump in reservoir activity $e^{\beta\mu}$, which is proportional to the jump in chemical potential $\mu$ when $\beta|\mu_\mathrm{i} - \mu_\mathrm{o}| \ll 1$.

For a charged ion, the energy landscape should include electrostatic energy in addition to the bare energetic contribution from the pore environment. Thus, the barrier and well energies become $\bar{E}_i^\ddagger = E_i^\ddagger + \mathrm{e}\phi_i^\ddagger$ and $\bar{E}_i = E_i + \mathrm{e}\phi_i$, where $\phi_i^\ddagger$ and $\phi_i$ denote the electric potentials at barrier $i$ and site $i$, respectively, and e is the fundamental charge. The same applies to the reservoirs, where the chemical potentials are now replaced by the electrochemical potentials $\bar{\mu} := \mu + \mathrm{e}\phi$.

#### (b). Equal barrier approximation

To further simplify the result, we use the idealized energy landscape shown in Fig. S2a. To that end, we consider sites with identical wells and barriers, where $E_i = E$ and $E_i^\ddagger = E + \mathcal{E}^\ddagger$, with $\mathcal{E}^\ddagger$ denoting the activation energy for each barrier crossing. In addition, equally spaced wells and barriers along the reaction coordinate yield a single local curvature scale at the energy extrema, whose square root yields the associated frequency factor, $\omega_i = \omega_i^\ddagger = \omega_0 \equiv \sqrt{\mathcal{E}^\ddagger}/(\delta/(2N))$ from Kramers' theory, where $\delta/N$ is the width of each energy barrier. We similarly specify the boundary frequency factor $\omega$ from the curvature scale associated with the entrance barrier. Using the barrier height $\mathcal{E}^\ddagger$ over the hydrated ion length scale $\ell_\mathrm{solv}$, the curvature scales as $\mathcal{E}^\ddagger/\ell_\mathrm{solv}^2$, leading to the frequency factor $\omega = \sqrt{\mathcal{E}^\ddagger}/\ell_\mathrm{solv}$.

The electrostatic contributions can be included through a linear electric potential across the membrane, which tilts the ionic energy landscape as shown in Fig. S2b. Choosing the outer membrane surface as the reference ($\phi_\mathrm{o} = 0$) and defining the membrane potential as $V \equiv \phi_\mathrm{i} - \phi_\mathrm{o} = \phi_\mathrm{i}$, the potentials at the barriers and wells are given by $\phi_i^\ddagger = V(2i-1)/(2N)$ and $\phi_i = Vi/N$, respectively. Incorporating the frequency factors and the electrostatic contributions into the ionic analogue of Eq. (S.19) and multiplying by the ion charge yields the steady-state electric current through the pore:

$$I_\mathrm{SS}^\mathrm{o} = \frac{\mathrm{e} D \beta \mathcal{E}^\ddagger e^{-\beta(E+\mathcal{E}^\ddagger)}}{\pi \delta \ell_\mathrm{solv}} \frac{e^{\beta\mu_\mathrm{o}} - e^{\beta(\mu_\mathrm{i} + \mathrm{e}V)}}{\frac{1}{N}\sum_{j=1}^{N} e^{\beta \mathrm{e}V(2j-1)/(2N)}} \ . \tag{S.20}$$

The subscript o denotes the steady-state current when the channel is open. When closed, the channel conducts no current.

The denominator has the form of a midpoint Riemann sum, so that in the limit of many barriers, $N \gg 1$,

$$\frac{1}{N}\sum_{j=1}^{N} e^{\beta \mathrm{e} V(2j-1)/(2N)} \;\approx\; \int_0^1 e^{\beta \mathrm{e} V x}\, dx \;=\; \frac{e^{\beta \mathrm{e} V}-1}{\beta \mathrm{e} V}\;. \tag{S.21}$$

For the idealized landscape with five barriers (Fig. S2c), the finite sum agrees well with the integral approximation: the relative error remains below $\sim 4\%$ for $0 \le \beta \mathrm{e}V \le 5$. We therefore replace the finite sum with its integral approximation in Eq. (S.21).

In the dilute-reservoir limit, we may further specify the reservoir chemical potentials as $\mu = \mu^{\ominus} + \beta^{-1}\ln\!\big(C_+/C^{\ominus}\big)$ [19]. Here, $C_+$ is the potassium concentration, $C^{\ominus}$ is the reservoir reference concentration, and $\mu^{\ominus}$ is the reservoir chemical potential at the reference concentration. Substituting these expressions into Eq. (S.20) yields

$$I^{\mathrm{o}}_{\mathrm{SS}} \;=\; \frac{\mathrm{e} D \beta \mathcal{E}^{\ddagger} e^{-\beta \mathcal{E}^{\ddagger}}}{\pi \delta \ell_{\mathrm{solv}} C^{\ominus} e^{\beta(E-\mu^{\ominus})}}\,\frac{\beta \mathrm{e} V\left(C^{\mathrm{o}}_+ - C^{\mathrm{i}}_+ e^{\beta \mathrm{e} V}\right)}{e^{\beta \mathrm{e} V}-1}\;, \tag{S.22}$$

where $C^{\mathrm{o}}_+$ and $C^{\mathrm{i}}_+$ are the outside and inside potassium concentrations, respectively. The remaining parameter to be discussed is the offset between the well energy and the reference chemical potential. This offset enters only through the factor $e^{\beta(E-\mu^{\ominus})}$, which plays the role of a partition coefficient between the reservoir and the internal sites. We therefore absorb it into an effective reference concentration in the pore, $C_{\mathrm{ref}} \equiv C^{\ominus} e^{\beta(E-\mu^{\ominus})}$, which is later determined by fitting to experimental data.

We note that the current law derived in Eq. (S.22) has the same functional form as the Goldman–Hodgkin–Katz flux equation [20, 21], where the analysis above allows us to identify an effective channel permeability as $P = D\beta\mathcal{E}^{\ddagger} e^{-\beta\mathcal{E}^{\ddagger}}/(\pi\delta\ell_{\mathrm{solv}} C_{\mathrm{ref}})$ in terms of microscopic parameters. Previous work by Hille [22] considered a similar energy-barrier and rate-theory perspective to derive Eq. (S.19), where all frequency factors are taken to be unity. Although Hille did not explicitly make the connection to the Goldman–Hodgkin–Katz flux equation, he did recognize that under certain conditions, multi-ion systems with this current relation may obey the Goldman–Hodgkin–Katz voltage equation.

#### (c). Symmetric reservoirs

In what follows, we specialize the constitutive relation to the case of initially symmetric reservoirs, with $C^{\mathrm{o}}_+ = C^{\mathrm{i}}_+ = C_0$. As shown in the next section, concentration perturbations arising from a transmembrane current are antisymmetric across the membrane. We may then define the concentration drop associated with a current $I^{\mathrm{o}}_{\mathrm{SS}}$ from outside to inside as $\Delta c_+ \equiv C^{\mathrm{i}}_+ - C^{\mathrm{o}}_+$, so that

$$C^{\mathrm{i}}_+ \;=\; C_0 + \Delta c_+/2\;, \qquad\qquad C^{\mathrm{o}}_+ \;=\; C_0 - \Delta c_+/2\;. \tag{S.23}$$

Substituting these expressions into Eq. (S.22), we obtain

$$I^{\mathrm{o}}_{\mathrm{SS}} \;=\; -\frac{\mathrm{e} D\beta\mathcal{E}^{\ddagger} e^{-\beta\mathcal{E}^{\ddagger}} C_0}{\pi\delta\ell_{\mathrm{solv}} C_{\mathrm{ref}}}\,\beta \mathrm{e} V\left[1 + \frac{\Delta c_+/(2C_0)}{\tanh(\beta \mathrm{e} V/2)}\right]\;. \tag{S.24}$$

Here, the prefactor multiplying the bracket represents the Ohmic current scale, while the terms inside the bracket account for corrections due to concentration perturbations. The Ohmic part then defines the single-channel conductance in the absence of concentration perturbations:

$$k \;=\; \frac{D\beta \mathrm{e}^2 C_0}{\pi\delta\ell_{\mathrm{solv}} C_{\mathrm{ref}}}\,\beta\mathcal{E}^{\ddagger} e^{-\beta\mathcal{E}^{\ddagger}}\;, \tag{S.25}$$

where $C_{\mathrm{ref}}$ is treated as a fitting parameter and is determined by comparison with experimental measurements. To account for gating, we restrict the current to the open state using the Heaviside function $\Theta(\xi)$, with $\Theta(\xi) = 1$ for $\xi \ge 0$ and $\Theta(\xi) = 0$ for $\xi < 0$, leading to the reduced form

$$I^{\mathrm{o}}_{\mathrm{SS}} \;=\; -\Theta(\xi) k V\left[1 + \frac{\Delta c_+/(2C_0)}{\tanh(\beta \mathrm{e} V/2)}\right]\;. \tag{S.26}$$

#### (d). Current relaxation to steady state

The steady-state current law in Eq. (S.26) assumes stationary pore occupancies. When the reservoir concentrations or gating state change, the relaxation dynamics of pore occupancies, and therefore the timescale to reach steady state, follow from the solution of the master equation (S.12), $\mathbf{x}(t) - \mathbf{x}_{\mathrm{SS}} = e^{-\mathbf{W}t}(\mathbf{x}(0) - \mathbf{x}_{\mathrm{SS}})$. The smallest eigenvalue of the rate matrix $\mathbf{W}$ in Eq. (S.13) thus determines the governing timescale for the current to reach its steady-state value.

Under the equal-barrier approximation introduced in the previous section, the rate matrix becomes

$$\mathbf{W} \;=\; \frac{2D}{\pi(\delta/N)^2}\beta\mathcal{E}^{\ddagger}e^{-\beta\mathcal{E}^{\ddagger}} \begin{bmatrix} e^{-\frac{\beta eV}{2N}} + e^{\frac{\beta eV}{2N}} & -e^{\frac{\beta eV}{2N}} & & \\ -e^{-\frac{\beta eV}{2N}} & e^{-\frac{\beta eV}{2N}} + e^{\frac{\beta eV}{2N}} & -e^{\frac{\beta eV}{2N}} & \\ & \ddots & & \\ & & -e^{-\frac{\beta eV}{2N}} & e^{-\frac{\beta eV}{2N}} + e^{\frac{\beta eV}{2N}} \end{bmatrix}, \tag{S.27}$$

which has a tridiagonal Toeplitz form. The eigenvalues of such matrices are well known [23] and are given by

$$\lambda_j \;=\; (4D\beta\mathcal{E}^{\ddagger}e^{-\beta\mathcal{E}^{\ddagger}}/(\pi(\delta/N)^2))[\cosh(\beta eV/(2N)) - \cos(j\pi/N)], \quad \text{with} \quad j \;=\; 1,\ldots,N-1\,. \tag{S.28}$$

The slowest relaxation mode corresponds to $j = 1$, yielding the governing relaxation timescale

$$\tau_1 \;=\; \lambda_1^{-1} \;=\; \frac{\pi(\delta/N)^2}{4D}\frac{e^{\beta\mathcal{E}^{\ddagger}}/(\beta\mathcal{E}^{\ddagger})}{\left[\cosh\left(\frac{\beta eV}{2N}\right) - \cos\left(\frac{\pi}{N}\right)\right]}\,, \tag{S.29}$$

which shows that relaxation becomes faster with a larger applied voltage and a lower barrier height.

As the number of barriers increases, this timescale saturates to

$$\tau_1 \;\sim\; \frac{\delta^2}{D}\frac{e^{\beta\mathcal{E}^{\ddagger}}}{\beta\mathcal{E}^{\ddagger}}\frac{2\pi}{(\beta eV)^2 + (2\pi)^2}\,. \tag{S.30}$$

For the idealized landscape with five barriers in Fig. S2c, the relaxation time is already close to this large $N$ limit over a physiologically relevant voltage range, $0 \le \beta e|V| \le 5$, with relative error below 5%. We therefore use Eq. (S.30) to estimate the longest relaxation time. Accordingly, we define the representative current relaxation timescale as the largest possible value of Eq. (S.30), obtained in the absence of a membrane potential:

$$\tau_{\mathrm{curr}} \;=\; \frac{\delta^2}{2\pi D}\frac{e^{\beta\mathcal{E}^{\ddagger}}}{\beta\mathcal{E}^{\ddagger}}\,, \tag{S.31}$$

which is the diffusive time needed to cross the membrane, increased by the energy barriers within the pore.

#### (e). Physiological values

We now estimate the current relaxation timescale $\tau_{\mathrm{curr}}$ and the reference concentration in the pore, $C_{\mathrm{ref}}$, using physiologically representative parameter values. For the relaxation time in Eq. (S.31), we use the membrane thickness $\delta \approx 4\,\mathrm{nm}$ and take the potassium diffusivity to be comparable to that in water, $D \approx 1\,\mathrm{nm}^2/\mathrm{ns}$, as estimated from molecular dynamics simulations [24]. We infer the barrier height $\mathcal{E}^{\ddagger}$ from computational studies of the energy profile through potassium channels. While these studies demonstrate that the positions of multiple ions in the pore can significantly affect the energy profile experienced by one ion due to ion–ion interactions [17], they provide an estimate of the energy-barrier scale, $\beta\mathcal{E}^{\ddagger} \approx 4$, at physiological temperature $T = 300$ K [17]. Together, these values give $\tau_{\mathrm{curr}} \sim 20\,\mathrm{ns}$ from Eq. (S.31). This value should be viewed as an upper-bound estimate of the relaxation time, since we use the full membrane thickness as the travel distance through the channel protein.

The conductance expression in Eq. (S.25) can also be used to infer the effective reference concentration in the pore, $C_{\mathrm{ref}} = C^{\ominus}e^{\beta(E-\mu^{\ominus})}$. Electrophysiological measurements report single-channel conductances for voltage-gated potassium channels in the range $k \sim 10$–$200\,\mathrm{pS}$ under physiological conditions [12, 25, 26]. Using the hydrated potassium length scale $\ell_{\mathrm{solv}} \approx 0.4\,\mathrm{nm}$ [27] and a typical physiological potassium concentration $C_0 \approx 150\,\mathrm{mM}$, we estimate $C_{\mathrm{ref}} \sim 70$–$1000\,\mathrm{mM}$, with larger values corresponding to smaller conductances. The inferred values of $C_{\mathrm{ref}}$ are comparable to physiological ionic concentrations, indicating that the site energy $E$ lies close to the chemical potential at physiological concentrations. This is consistent with the established principle of potassium selectivity: the binding sites in the selectivity filter compensate the desolvation cost by mimicking key aspects of the potassium hydration environment [15].

Lastly, Eq. (S.26) gives the steady-state behavior of the channel current for both open and closed states. Two processes control the transition between the open and closed states. First, the channel continuously changes conformations between the open and closed states, which may be conductive or partially conductive. Moreover, the energy landscape for an ion within the channel, approximated in Fig. S2, dynamically evolves during this process. The timescale to commit to one conformation from the transition state can be approximated by $\zeta/U^{\ddagger} \sim 10\,\mathrm{ns}$, i.e., the ratio of the channel friction coefficient to the energy barrier. Once the channel has committed to either conformation, $\tau_{\mathrm{curr}}$ governs the adjustment of the current through the channel. Therefore, upon transitioning from closed to open, we expect the channel to reach a steady-state current on a timescale set by the sum of these two timescales, which remains of order $\tau_{\mathrm{curr}}$. From Sec. I.1 (a), we estimate the gating timescale $\tau_{\mathrm{G}} \sim 40\,\mu\mathrm{s} \gg \tau_{\mathrm{curr}}$. Therefore, we can neglect these fast relaxation processes and take the instantaneous current to follow the steady-state current, such that

$$I(t) = -\Theta(\xi(t))kV(t)\left[1 + \frac{\Delta c_{+}(t)/(2C_0)}{\tanh(\beta eV(t)/2)}\right], \quad \text{(S.32)}$$

which is the relation described in the main text.

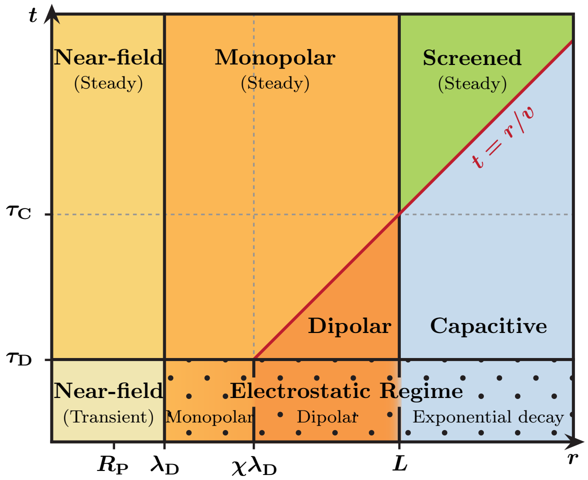

Figure S3: Given a constant current $I$ through an ion channel flanked by two Faradaic electrodes at a distance $L$, the diagram depicts the different regimes of the membrane potential's response as a function of distance $r$ from the channel and time $t$ from channel opening. Electrostatic effects dominate for $t < \tau_{\mathrm{D}}$ as charge reorganization occurs on timescales of $\tau_{\mathrm{D}}$ and larger. Beyond this, a steady-state emerges for $t > r/v$ at any position $r$, where $v$ is the propagation speed. This steady state is monopolar ($1/r$) for $r < L$ and exponential ($\exp(-\pi r/2L)$) for $r > L$.

#### 3. Effects of single channel current

A localized transmembrane current through a single ion channel perturbs both the electric potential and ion concentrations. Here, we summarize the boundary layer analysis of the resulting electrochemical response in Ref. [28] for a membrane placed between Faradaic electrodes, with the corresponding spatiotemporal regimes reproduced in Fig. S3.

We consider an impermeable membrane of thickness $\delta$ and dielectric permittivity $\epsilon \sim 4\epsilon_0$ separating two symmetric reservoirs of a binary monovalent salt with bulk concentration $C_0$ ($\sim 150\,\mathrm{mM}$). Faradaic plate electrodes are placed parallel to the membrane at a distance $L$ from each membrane surface (as in Fig. 1a of the main text) and are held at zero electric potential. At $t = 0$, a constant cationic current $I$ is imposed through a circular pore of radius $R_{\mathrm{P}}$ from outside to inside the membrane, mimicking the opening of a single channel.

Charges transported through the membrane reorganize within diffuse charge layers near the membrane surfaces. The thickness of these diffuse layers is characterized by the Debye length $\lambda_{\mathrm{D}}$ ($\sim 1\,\mathrm{nm}$). The associated Debye time $\tau_{\mathrm{D}} = \lambda_{\mathrm{D}}^2/D$ ($\sim 1\,\mathrm{ns}$) defines the diffusive time for local charge relaxation. This nanoscale charge reorganization underlies the macroscopic spreading of the electric potential and charge density along the membrane, resulting in the emergent spatiotemporal regimes depicted in Fig. S3.

We focus primarily on the membrane potential $V(r,t)$, the charge density at the membrane surface $\rho(r,t)$, and the salt concentration perturbation at the membrane surface $C(r,t)$, where $r$ is the distance along the membrane from the center of the pore. Consistent with the definition in the preceding section, the membrane potential $V = \phi_\mathrm{i} - \phi_\mathrm{o}$ is referenced to the outside. The charge and salt concentration perturbations are also evaluated on the inside surface, so that the current-induced perturbations $V$, $\rho$, and $C$ have the same sign as the current. By symmetry, the charge and concentration perturbations on the outside surface have the opposite sign. We define the charge density as $\rho = \mathrm{e}(C_+ - C_-)$ and the salt concentration perturbation as $C = C_+ + C_- - 2C_0$, where $C_+$ and $C_-$ are the cation and anion concentrations on the inside membrane surface. The cation concentration jump across the membrane, defined in Eq. (S.23), is then written in terms of the charge density and salt concentration perturbation as

$$\Delta c_+ \; = \; 2\,(C_+ - C_0) \; = \; C + \frac{\rho}{\mathrm{e}} \; . \tag{S.33}$$

The electrochemical response of the system subjected to a localized current can be analyzed using the Poisson-Nernst-Planck framework, as detailed in Refs. [28–30]. The full Poisson-Nernst-Planck equations governing the coupled evolution of the membrane potential, charge density, and salt concentration are nonlinear. However, as argued in Ref. [28], the current through an individual channel is sufficiently small under physiological conditions that the linearized dynamics accurately captures the electrochemical response. In that case, the problem can be formulated in terms of Green's functions [29], with each field scaling linearly with the imposed current $I$:

$$V(r,t) \; = \; I\int_0^t G_\mathrm{V}(r,t-\tau)\,d\tau \; , \quad \rho(r,t) \; = \; I\int_0^t G_\rho(r,t-\tau)\,d\tau \; , \quad C(r,t) \; = \; I\int_0^t G_\mathrm{C}(r,t-\tau)\,d\tau \; . \tag{S.34}$$

Here, $G_\mathrm{V}$, $G_\rho$, and $G_\mathrm{C}$ are the Green's functions describing the respective responses of the membrane potential, charge density, and salt concentration to a unit impulse of channel current, i.e., an imposed current through the pore $I(t) = \delta(t)$. More generally, this linearity allows the electrochemical response to an arbitrary time-dependent channel current $I(t)$ to be written in terms of the Green's functions as

$$V(r,t) \; = \; \int_{-\infty}^t I(\tau) G_\mathrm{V}(r,t-\tau)\,d\tau \; , \tag{S.35}$$

$$\rho(r,t) \; = \; \int_{-\infty}^t I(\tau) G_\rho(r,t-\tau)\,d\tau \; , \tag{S.36}$$

$$C(r,t) \; = \; \int_{-\infty}^t I(\tau) G_\mathrm{C}(r,t-\tau)\,d\tau \; . \tag{S.37}$$

The Green's functions are obtained from the step-current solutions in Ref. [28], using the relation between a step response and an impulse response in Eq. (S.34). After normalization by the imposed current, differentiating each step-current response with respect to time gives the corresponding Green's function.

The electrochemical response to step currents can be classified into three regimes [28]: (i) the near-field regime ($r < R_\mathrm{P}$), (ii) the intermediate regime ($\lambda_\mathrm{D} < r < L$), and (iii) the far-field regime ($r > L$). We first consider the near-field regime and discus the self-response at the channel location, $r = 0$, for a pore radius $R_\mathrm{P}$. From the near-field response to a step current [28, 29], we obtain

$$G_\mathrm{V}(0,t) \; = \; \frac{2}{\pi R_\mathrm{P}^2 \epsilon/\lambda_\mathrm{D}} e^{-\bar{t}} \left[ \bar{R}_\mathrm{P}\,\mathrm{erf}\left(\frac{\bar{R}_\mathrm{P}}{2\sqrt{\bar{t}}}\right) - \frac{2\sqrt{\bar{t}}}{\sqrt{\pi}} \left(1 - e^{-\bar{R}_\mathrm{P}^2/(4\bar{t})}\right)\right] \; , \tag{S.38}$$

$$G_\rho(0,t) \; = \; \frac{1}{\pi R_\mathrm{P}^2 \lambda_\mathrm{D}} \frac{e^{-\bar{t}}}{\sqrt{\pi \bar{t}}} \left[1 - e^{-\bar{R}_\mathrm{P}^2/(4\bar{t})}\right] \; , \tag{S.39}$$

$$G_\mathrm{C}(0,t) \; = \; \frac{1}{\pi R_\mathrm{P}^2 \lambda_\mathrm{D} \mathrm{e}} \frac{1}{\sqrt{\pi \bar{t}}} \left[1 - e^{-\bar{R}_\mathrm{P}^2/(4\bar{t})}\right] \; . \tag{S.40}$$

where $\bar{t} = t/\tau_\mathrm{D}$ and $\bar{R}_\mathrm{P} = R_\mathrm{P}/\lambda_\mathrm{D}$.

For intermediate distances from the pore, larger than the Debye length but smaller than the electrode separation, $\lambda_\mathrm{D} < r < L$, and for times longer than the Debye time, $t > \tau_\mathrm{D}$ (Fig. S3), electrode effects are negligible and the electrochemical signal propagates ballistically along the membrane [28, 29]. The corresponding

Green's functions are

$$G_{\rm V}(r,t) = \frac{1}{g}\frac{\chi}{1+\chi}\frac{v^2 t}{\pi\left(r^2+(vt)^2\right)^{3/2}} , \tag{S.41}$$

$$G_{\rho}(r,t) = \frac{1}{D}\frac{1}{1+\chi}\frac{v^2 t}{2\pi\left(r^2+(vt)^2\right)^{3/2}} , \tag{S.42}$$

$$G_{\rm C}(r,t) = \frac{2}{\rm e}\frac{e^{-r^2/(4Dt)}}{(4\pi Dt)^{3/2}} . \tag{S.43}$$

where $\chi = (\epsilon/(2\lambda_{\rm D}))/(\epsilon^{\rm M}/\delta) \gg 1$ is the capacitance ratio between the electrical double layers and the membrane, $v = (1+\chi)\lambda_{\rm D}/\tau_{\rm D}$ is the signal propagation speed, and $g = D\epsilon/\lambda_{\rm D}^2$ is the electrolyte conductivity. The spatiotemporal structure of $G_{\rm V}$ and $G_\rho$ is controlled by the factor $t/(r^2+(vt)^2)^{3/2}$, which encodes electrochemical signal propagation in the intermediate region $\lambda_{\rm D} < r < L$. For $vt \ll r$, this factor scales as $t/r^3$, corresponding to the early dipolar response. Once the signal reaches the distance $r$, i.e. $vt \gtrsim r$, the response crosses over toward the intermediate monopolar regime, which gives the steady monopolar scaling $1/r$ under a constant current, as shown in Fig. S3. Note that the salt concentration perturbation decouples from the electrostatic fields under the linearized dynamics and follows the diffusion equation. Accordingly, $G_{\rm C}$ in Eq. (S.43) has the form of the ordinary three-dimensional diffusion Green's function. We note that this form remains valid for times shorter than the *salt diffusion timescale* across the electrode separation, given by $\tau_{\rm L} = L^2/D$. At longer times, $t \gg \tau_{\rm L}$, the salt concentration dynamics effectively become two-dimensional due to the slab geometry of the electrolyte, and the Faradaic reactions at the electrodes may also affect the salt response near the membrane.

In the far-field regime, $r > L$, the salt concentration perturbation remains the same as in the intermediate regime, given by Eq. (S.43). In contrast, the far-field Green's functions for the membrane potential and charge density differ from those in the intermediate regime:

$$G_{\rm V}(r,t) = \frac{2\chi}{1+\chi}\sqrt{\frac{\pi t L}{r\tau_{\rm C}}}K_0\left(\frac{\pi r}{L}\right)e^{-t/\tau_{\rm C}}I_1\left(2\sqrt{\frac{\pi r t}{L\tau_{\rm C}}}\right) , \tag{S.44}$$

$$G_{\rho}(r,t) = \frac{1}{1+\chi}\sqrt{\frac{\pi t L}{r\tau_{\rm C}}}K_0\left(\frac{\pi r}{L}\right)e^{-t/\tau_{\rm C}}I_1\left(2\sqrt{\frac{\pi r t}{L\tau_{\rm C}}}\right) , \tag{S.45}$$

which show that electrode screening produces exponential decay in space over the electrode separation distance $L$ and in time over the membrane capacitive timescale [30, 31],

$$\tau_{\rm C} = \frac{L\lambda_{\rm D}}{D(1+\chi)} = \frac{L}{v} . \tag{S.46}$$

The influence of these screened fields arising from the single-channel response can be neglected when considering the multi-channel system.

#### (a). Approximation by exponential functions

The Green's functions encode memory in the electrochemical response: the membrane potential, charge density, and concentration fields at a given time depend on the history of channel currents. The memory encoded by the Green's functions in Eqs. (S.38)–(S.42) decays over the characteristic timescales $\tau_{\rm D}$, $\tau_{\rm ch}$, and $r/v$, each of which is significantly shorter than the gating timescale $\tau_{\rm G}$. Thus, once a channel opens, the electrochemical response and the resulting current $I(t)$ quickly relax and remain at their steady-state values over most of the time that the open state is maintained.

This separation of timescales implies that the dominant contribution of the memory kernels to the electrochemical dynamics is determined by their response under a sustained current. Thus, the Green's functions can be approximated by kernels that preserve both the characteristic relaxation time and the response to a constant current. Although this approximation does not resolve the detailed transient electrochemical response over the short memory timescales, it remains accurate on the timescales of interest, where channels undergo multiple gating events, to leading order in the ratio of the memory timescales to $\tau_{\rm G}$. This provides a significant computational advantage when evaluating the convolution integrals in Eqs. (S.35)–(S.37).

Direct evaluation of the convolution integrals with the exact Green's functions in Eqs. (S.38)–(S.42) is computationally prohibitive, since the time integral needs to be recomputed over the full current history at each time point. However, if one were to approximate the exact Green's functions, as mentioned above, by exponentially decaying functions in time, the integrals can be computed recursively without retaining the full time history, as discussed in Sec. III. One must therefore find suitable exponential approximations for each true Green's function. To this end, we approximate each Green's function as an exponential function of the form $A(r)e^{-t/T(r)}$, where the amplitude $A(r)$ and timescale $T(r)$ may depend on the distance from the pore, $r$. The timescale $T(r)$ must be chosen to represent the characteristic relaxation time of the original Green's function. The amplitude $A(r)$ is then determined by requiring the approximation and the exact kernel to have the same time integral, thereby preserving the steady-state response to a constant current.

For the self-response at the channel location $r = 0$, we use the leading-order approximation in the limit $R_{\rm P}/\lambda_{\rm D} \ll 1$. The resulting exponential kernels, whose time integrals match those of the corresponding exact Green's functions, are

$$\tilde{G}_{\rm V}(0,t) = \frac{1}{\pi g \lambda_{\rm D}} \frac{e^{-t/\tau_{\rm D}}}{\tau_{\rm D}} , \qquad \tilde{G}_{\rho}(0,t) = \frac{1}{\pi D R_{\rm P}} \frac{e^{-t/\tau_{\rm ch}}}{\tau_{\rm ch}} , \qquad \tilde{G}_{\rm C}(0,t) = \frac{1}{\pi D R_{\rm P} {\rm e}} \frac{e^{-t/\tau_{\rm ch}}}{\tau_{\rm ch}} . \tag{S.47}$$

Thus, while the self-response of the membrane potential is characterized by the Debye relaxation time $\tau_{\rm D}$, the charge density and salt concentration are characterized by the diffusive timescale over the pore size, $\tau_{\rm ch} = R_{\rm P}^2/D$.

For the nonlocal response, we first consider the intermediate region $\lambda_{\rm D} < r < L$, where electrode screening is not yet dominant. The exponential kernels that preserve the associated timescales and time integrals are given by

$$\tilde{G}_{\rm V}(r,t) = \frac{\chi}{1+\chi} \frac{1}{\pi g r} \frac{e^{-t/\tau_{\rm V}(r)}}{\tau_{\rm V}(r)} , \tag{S.48}$$

$$\tilde{G}_{\rho}(r,t) = \frac{1}{1+\chi} \frac{1}{2\pi D r} \frac{e^{-t/\tau_{\rm V}(r)}}{\tau_{\rm V}(r)} , \tag{S.49}$$

$$\tilde{G}_{\rm C}(r,t) = \frac{1}{2\pi D r {\rm e}} \frac{e^{-t/\tau_{\rm diff}(r)}}{\tau_{\rm diff}(r)} . \tag{S.50}$$

Here, the salt concentration response is characterized by the diffusive timescale $\tau_{\rm diff}(r) = \pi^2 r^2/(4D)$. In contrast, the membrane potential and charge density responses are characterized by $\tau_{\rm V}(r)$, the timescale required for the electrical responses in Eqs. (S.35) and (S.36) to reach the steady monopolar regime under a constant current, which is illustrated in Fig. S3. Close to the pore, $r \lesssim v\tau_{\rm D}$, this occurs within the Debye relaxation time $\tau_{\rm D}$ upon formation of the diffuse charge layer. More distant regions reach this regime only after the electrochemical signal arrives, which takes time $r/v$. Thus, for the membrane potential and charge density kernels, the signal propagation timescale $\tau_{\rm V}(r) = \max(\tau_{\rm D}, r/v)$ characterizes the crossover to the monopolar regime from the dipolar regime, under a sustained current.

For distances larger than the electrode separation, $r > L$, we continue to use the same exponential approximation $\tilde{G}_{\rm C}(r,t)$ in Eq. (S.50) for the salt concentration, since the nonlocal salt concentration response does not distinguish whether $r$ is smaller or larger than $L$. In contrast, the membrane potential and charge density responses are exponentially small due to electrode screening, scaling as $e^{-\pi r/(2L)}$. We therefore neglect these kernels for $r > L$, i.e., $\tilde{G}_{\rm V}(r,t) = \tilde{G}_{\rho}(r,t) = 0$.

#### (b). Applied electrode potential

In the preceding discussions, the electrodes bounding the membrane are held at zero potential, and the electrochemical response is driven only by the transmembrane current through a channel. We now consider the electrochemical response when a nonzero electrode potential $V_{\rm ext}$ is also applied between the two electrodes. In the linear regime considered here, responses to imposing an additional electrode potential amount to a superposition of the *current-driven* response, where the electrodes are held at zero potential, and the *voltage-driven* response, where no transmembrane current is imposed but the electrodes are held at $V_{\rm ext}$, as shown in Ref. [28]. The *voltage-driven* response has been studied in Refs. [30, 31] and is summarized as follows.

When the Faradaic electrodes impose the electrode potential difference $V_{\rm ext}$, defined as the inside electrode potential relative to the outside electrode potential, the transmembrane potential reaches the steady value $[\chi/(\chi+1)]V_{\rm ext}$ on the membrane capacitive timescale $\tau_{\rm C}$, defined in Eq. (S.46) [30, 31]. Since $\chi \gg 1$, $[\chi/(\chi+1)]V_{\rm ext} \approx V_{\rm ext}$, indicating that much of the electrode potential difference is borne by the membrane. Accordingly, the charge density at steady state is given by the local capacitance of the membrane as $\rho = gV_{\rm ext}/(2(\chi+1)D)$, while the salt concentration is unaffected to linear order.

With both a transmembrane current and an applied electrode potential, we thus have

$$V(r,t) \;=\; \frac{\chi}{\chi+1}V_{\rm ext} + \int_{-\infty}^{t} I(\tau)G_{\rm V}(r,t-\tau)\,d\tau \;, \tag{S.51}$$

$$\rho(r,t) \;=\; \frac{g}{2(\chi+1)D}V_{\rm ext} + \int_{-\infty}^{t} I(\tau)G_{\rho}(r,t-\tau)\,d\tau \;, \tag{S.52}$$

$$C(r,t) \;=\; \int_{-\infty}^{t} I(\tau)G_{\rm C}(r,t-\tau)\,d\tau \;. \tag{S.53}$$

We neglect the transient dynamics induced by the applied potential over the membrane capacitive timescale $\tau_{\rm C}$, not because this timescale is necessarily shorter than the others, but because we focus on the steady state reached after sustained maintenance of the external electrode potential.

#### 4. Single-channel response and parameter estimation

The results and simplifications above allow us to study the response of a single channel under an applied voltage. To that end, the governing equations for the single-channel stochastic dynamics are the overdamped Langevin equation (S.3) for channel gating, the instantaneous current law in Eq. (S.32), and the exponential memory kernels for the self-response in Eq. (S.47):

$$\zeta\frac{d\xi(t)}{dt} \;=\; -4U^{\ddagger}\left(\xi(t)^3-\xi(t)\right) + Q\left(V(t)-V_{\rm G}\right) + \eta(t) \;, \tag{S.54}$$

$$\langle\eta(t)\rangle \;=\; 0 \;, \qquad \langle\eta(t)\eta(t')\rangle \;=\; 2\zeta k_{\rm B}T\,\delta(t-t') \;, \tag{S.55}$$

$$I(t) \;=\; -\Theta\left(\xi(t)\right)kV(t)\left[1+\frac{1}{2C_0}\frac{C(t)+\rho(t)/{\rm e}}{\tanh\left(\beta {\rm e}V(t)/2\right)}\right] \;, \tag{S.56}$$

$$V(t) \;=\; \frac{\chi}{\chi+1}V_{\rm ext} + \frac{1}{\pi g\lambda_{\rm D}}\int_0^t I(\tau)\frac{e^{-(t-\tau)/\tau_{\rm D}}}{\tau_{\rm D}}\,d\tau \;, \tag{S.57}$$

$$\rho(t) \;=\; \frac{g}{2(\chi+1)D}V_{\rm ext} + \frac{1}{\pi DR_{\rm P}}\int_0^t I(\tau)\frac{e^{-(t-\tau)/\tau_{\rm ch}}}{\tau_{\rm ch}}\,d\tau \;, \tag{S.58}$$

$$C(t) \;=\; \frac{1}{\pi DR_{\rm P}{\rm e}}\int_0^t I(\tau)\frac{e^{-(t-\tau)/\tau_{\rm ch}}}{\tau_{\rm ch}}\,d\tau \;, \tag{S.59}$$

where $V(t)$, $\rho(t)$, and $C(t)$ denote the local membrane potential, charge density, and salt concentration perturbation at the channel position. Although the only stochastic equation is the Langevin equation for the gating variable $\xi$, the resulting stochasticity propagates through the current law and the convolution integrals. The numerical implementation of the stochastic differential equation and convolution integrals is described in Sec. III.

##### (a). Limiting behaviors of current

Since the channel remains open (on average) much longer than the Debye time, i.e. $\tau_{\rm o\to c} \gg \tau_{\rm D}$, the self-response induced by a current reaches steady state while the channel is open. We therefore use the steady-state values of the current, local membrane potential, charge density, and concentration perturbation. The resulting self-consistent steady-state current is described by

$$I \;=\; -k\left(\frac{\chi}{\chi+1}V_{\rm ext}+\frac{I}{\pi g\lambda_{\rm D}}\right)\left[1+\frac{\frac{g}{4C_0D{\rm e}}\left(\frac{V_{\rm ext}}{\chi+1}+\frac{4I}{\pi gR_{\rm P}}\right)}{\tanh\left(\frac{\beta{\rm e}}{2}\left(\frac{\chi}{\chi+1}V_{\rm ext}+\frac{I}{\pi g\lambda_{\rm D}}\right)\right)}\right] \;, \tag{S.60}$$

which gives an implicit current-voltage relation for a single channel.

For a dilute electrolyte, the conductivity, diffusivity, and temperature are related by $g/(4C_0 D\mathrm{e}) = \beta\mathrm{e}/2$. As discussed in the following section, we neglect the temperature dependence of the parameters, so, to maintain self-consistency with this relation, we define a reference temperature by $\beta_{\mathrm{ref}} = g/(2C_0 D\mathrm{e}^2)$ and take $\beta = \beta_{\mathrm{ref}}$ in the current-voltage relation in Eq. (S.60). We nondimensionalize this constitutive relation by the dimensionless current $\tilde{I} = I/(k[\chi/(\chi+1)]V_{\mathrm{ext}})$ and dimensionless applied potential $\hat{V}_{\mathrm{ext}} = \beta_{\mathrm{ref}}\mathrm{e}[\chi/(\chi+1)]V_{\mathrm{ext}}/2$. This gives

$$\tilde{I} \;=\; -(1+2\tilde{\gamma}\tilde{I})\left(1+\frac{\hat{V}_{\mathrm{ext}}\left(\frac{1}{\chi}+8\tilde{\gamma}\frac{\lambda_{\mathrm{D}}}{D}\tilde{I}\right)}{\tanh\left(\hat{V}_{\mathrm{ext}}(1+2\tilde{\gamma}\tilde{I})\right)}\right) \;, \tag{S.61}$$

where $\tilde{\gamma} = k/(2\pi g\lambda_{\mathrm{D}}) \approx 4.6\times 10^{-2}$ characterizes the strength of the self-response in the local membrane potential for a given externally imposed potential. Physically, $\tilde{\gamma}$ compares the channel conductance $k$ with the access-like conductance of the electrolyte dome of radius $\lambda_{\mathrm{D}}$ around the channel, where the electrical self-response occurs: $g(2\pi\lambda_{\mathrm{D}}^2)/\lambda_{\mathrm{D}}$, with area $2\pi\lambda_{\mathrm{D}}^2$ and length $\lambda_{\mathrm{D}}$. Larger $\tilde{\gamma}$ indicates that the electrolyte less effectively relaxes the charges supplied by the current through the channel, leading to a stronger self-response in the local membrane potential.

We consider two limiting scenarios of the applied potential, one in which it is small and one in which it is large. For $|\hat{V}_{\mathrm{ext}}| \ll 1$, we expand the governing equation in a series about $\hat{V}_{\mathrm{ext}} = 0$ and solve for the current $\tilde{I}$ in powers of $\hat{V}_{\mathrm{ext}}$. This gives, to leading order,

$$\tilde{I} \;=\; -\frac{\chi+1}{\chi}\frac{1}{1+2\tilde{\gamma}\left(1+4\frac{\lambda_{\mathrm{D}}}{R_{\mathrm{P}}}\right)} + \mathcal{O}\left(\hat{V}_{\mathrm{ext}}^2\right) \;, \tag{S.62}$$

where the odd powers in $\hat{V}_{\mathrm{ext}}$ vanish since $\tilde{I}$ is even in $\hat{V}_{\mathrm{ext}}$. Restoring dimensions, this result gives

$$I \;=\; -\frac{V_{\mathrm{ext}}}{\frac{1}{k}+\frac{1}{\pi g\lambda_{\mathrm{D}}}+\frac{4}{\pi g R_{\mathrm{P}}}} \;. \tag{S.63}$$

We can identify an effective channel conductance as

$$\frac{1}{k_{\mathrm{eff}}} \;=\; \frac{1}{k}+\frac{1}{\pi g\lambda_{\mathrm{D}}}+\frac{4}{\pi g R_{\mathrm{P}}} \;. \tag{S.64}$$

Here, the second and third terms on the right-hand side represent the self-responses from the membrane potential and concentration jump, respectively. The linear relation in Eq. (S.63) is now written simply as $I = -k_{\mathrm{eff}}V_{\mathrm{ext}}$, which is the conventional linear current-voltage relation used to define the channel conductance from experimental measurements.

For large applied potentials, $|\hat{V}_{\mathrm{ext}}| \gg 1$, the asymptotic behavior $\tanh(x) \to \mathrm{sgn}(x)$ as $|x| \to \infty$ reduces the implicit current law in Eq. (S.60) to

$$\tilde{I} \;=\; -(1+2\tilde{\gamma}\tilde{I})\left[1+|\hat{V}_{\mathrm{ext}}|\left(\frac{1}{\chi}+8\tilde{\gamma}\frac{\lambda_{\mathrm{D}}}{R_{\mathrm{P}}}\tilde{I}\right)\right] \;. \tag{S.65}$$

We then expand $\tilde{I}$ in inverse powers of $|\hat{V}_{\mathrm{ext}}|$ to obtain

$$\tilde{I} \;=\; -\frac{1}{8\chi\frac{\lambda_{\mathrm{D}}}{R_{\mathrm{P}}}\tilde{\gamma}}\left[1+\frac{\chi}{|\hat{V}_{\mathrm{ext}}|}\left(1-\frac{1}{2\tilde{\gamma}\left(4\chi\frac{\lambda_{\mathrm{D}}}{R_{\mathrm{P}}}-1\right)}\right)+\mathcal{O}\left(|\hat{V}_{\mathrm{ext}}|^{-2}\right)\right] \;. \tag{S.66}$$

Here, we retain the first-order correction, since it becomes a constant voltage offset in dimensional form:

$$I \;=\; -k_{\infty}\mathrm{sgn}(V_{\mathrm{ext}})\left(|V_{\mathrm{ext}}|+V_{\mathrm{res}}\right) \;, \tag{S.67}$$

where

$$k_{\infty} \;=\; \frac{\pi g R_{\mathrm{P}}}{4(\chi+1)} \;, \qquad V_{\mathrm{res}} \;=\; \frac{2(\chi+1)}{\beta_{\mathrm{ref}}\mathrm{e}}\left(1-\frac{\frac{1}{k}}{\frac{\chi}{\chi+1}\frac{1}{k_{\infty}}-\frac{1}{\pi g\lambda_{\mathrm{D}}}}\right) \;. \tag{S.68}$$

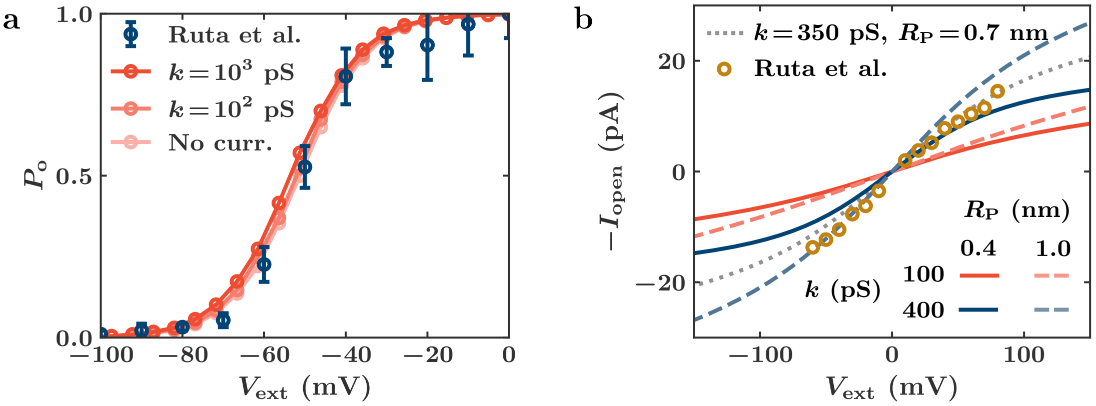

Figure S4: (a) The probability that a single channel is open as a function of the applied potential $V_{\text{ext}}$. The experimental data are from Ref. [12]. The lightest curve shows the behavior with no current ($k = 0$), while the other two correspond to behaviors at large channel conductances. Even beyond physiologically relevant values of $k$, the response curve of a single channel is relatively insensitive, indicating that the self-response has little effect on the steady-state open probability. Both curves that include current use $R_{\text{P}} = 0.4\,\text{nm}$. (b) The dependence of the open current $I_{\text{open}}$ on the applied voltage $V_{\text{ext}}$ for a single channel, as determined by Eq. (S.60). The experimental data are from Ref. [12]. The dotted line shows the behavior for the fitted values $k = 350\,\text{pS}$ and $R_{\text{P}} = 0.7\,\text{nm}$. The other curves show how the response changes with $k$ and $R_{\text{P}}$.

The current law again becomes linear in the applied voltage in the large-potential limit, but with a different effective conductance $k_\infty \approx 16\,\text{pS}$ and a residual voltage offset $V_{\text{res}} \approx 2\,\text{V}$. The effective conductance $k_\infty$ is the conductance of the electrolyte region over the pore radius, where the concentration response takes place, and is reduced by the factor $\chi + 1 \gg 1$. This reduction reflects the fact that the concentration response in the electrolyte is driven by a much smaller effective potential scale than the membrane potential. The independence of $k_\infty$ from the channel conductance $k$ shows that the concentration correction, rather than the intrinsic channel resistance, becomes the main bottleneck for the current when the applied potential is large.

The effect of intrinsic channel conductance enters only through the next-order correction in $V_{\text{res}}$. Although $V_{\text{res}}$ becomes negligible compared with $|V_{\text{ext}}|$ as $|\hat{V}_{\text{ext}}| \to \infty$, it provides a meaningful correction at physiologically accessible large potentials due to the enhancement of its scale by $\chi + 1$.

#### (b). Model parameters

The full set of parameters in the single-channel model consists of the drag coefficient for the gating variable $\zeta$, the energy barrier for the gating transition $U^\ddagger$, the effective gating charge $Q$, the gating voltage $V_{\text{G}}$, the temperature $T$, the single-channel conductance $k$, the initial bulk concentration $C_0$, the externally applied potential $V_{\text{ext}}$, the electrolyte conductivity $g$, the Debye length $\lambda_{\text{D}}$, the Debye timescale $\tau_{\text{D}}$, the ionic diffusivity $D$, the effective pore radius $R_{\text{P}}$, the pore diffusion timescale $\tau_{\text{ch}}$, and the capacitance ratio $\chi$. Among these, the electrolyte conductivity, the Debye timescale, and the pore diffusion timescale are determined by the other parameters via $g = 2\beta e^2 C_0 D$, $\tau_{\text{D}} = \lambda_{\text{D}}^2/D$, and $\tau_{\text{ch}} = R_{\text{P}}^2/D$, respectively. Thus, the model contains 12 independent parameters, and we freely vary $V_{\text{ext}}$ and $T$ as control parameters.

The remaining 10 independent parameters are grouped as follows: (i) $\zeta$, $U^\ddagger$, $Q$, and $V_{\text{G}}$ are the gating parameters, (ii) $C_0$, $\lambda_{\text{D}}$, and $D$ are the electrolyte parameters, (iii) $\chi$ is the membrane parameter, and (iv) $k$ and $R_{\text{P}}$ are the current parameters. We use $\zeta = 2.5\,\text{eV}\cdot\text{ns}$ and $U^\ddagger = 0.2\,\text{eV}$, obtained by fitting to physiological measurements as described in Sec. I.1 (a). We also take a physiologically representative concentration $C_0 = 150\,\text{mM}$, which gives a Debye length $\lambda_{\text{D}} = 1\,\text{nm}$ at room temperature. The potassium ion diffusivity is taken as $D = 1\,\text{nm}^2/\text{ns}$, and the capacitance ratio between the electrical double layers and the membrane is set to $\chi = 40$, a value typical of physiological conditions [28–30].

This leaves four parameters to be determined: the gating parameters $Q$ and $V_{\text{G}}$ and the current pa-

| Parameter | Symbol | Value |
|---|---|---|
| Gate charge | $Q$ | 1.6e |
| Gating voltage | $V_{\mathrm{G}}$ | 50 mV |
| Gating variable drag | $\zeta$ | 2.5 eV · ns |
| Transition barrier | $U^{\ddagger}$ | 0.2 eV |
| Initial concentration | $C_0$ | 150 mM |
| Debye length | $\lambda_{\mathrm{D}}$ | 1 nm |
| Ionic diffusivity | $D$ | 1 nm$^2$/ns |
| Electrolyte conductivity | $g$ | 1.2 S/m |
| Debye time | $\tau_{\mathrm{D}}$ | 1 ns |
| Capacitance ratio | $\chi$ | 40 |
| Channel conductance | $k$ | 350 pS |
| Pore radius | $R_{\mathrm{P}}$ | 0.7 nm |
| Pore diffusion time | $\tau_{\mathrm{ch}}$ | 0.09 nm$^2$/ns |

Table S1: Parameters for single-channel simulations. For multiple-channel simulations, we retain all these parameters, with additional parameters capturing the channel-to-channel distance and the electrode separation.

rameters $k$ and $R_{\mathrm{P}}$. We estimate these using single-channel open probability and single-channel current data for the KvAP channel in Ref. [12]; this channel is also the focus of the main text. We first determine the gating parameters $Q$ and $V_{\mathrm{G}}$ from the open probability data using Eq. (S.8). In the fitting, we neglect the self-response induced by the current and use $V = V_{\mathrm{ext}}$. This gives $Q \approx 1.6\mathrm{e}$ and $V_{\mathrm{G}} = -50\,\mathrm{mV}$, close to the values reported in Ref. [12] by fitting the same Boltzmann form in Eq. (S.8). Including the current self-response slightly perturbs the open probability, as shown in Fig. S4a, but the differences remain within the uncertainty of the experimental measurements.

We determine $k$ and $R_{\mathrm{P}}$ by fitting the implicit current-voltage relation in Eq. (S.60) to the single-channel current data in Ref. [12], as shown in Fig. 1f of the main text and Fig. S4b. The current magnitude increases with both the channel conductance $k$ and the pore radius $R_{\mathrm{P}}$. The pore radius affects the current through the concentration jump induced by the current—a larger pore radius reduces this concentration jump and therefore weakens the self-response opposing the current. The implicit relation also exhibits nonlinear behavior at large applied voltages. The nonlinear deviation begins at similar values of $V_{\mathrm{ext}} \sim \pm 50\,\mathrm{mV}$ (corresponding to $\hat{V}_{\mathrm{ext}} \sim 1$) for all values of $k$ and $R_{\mathrm{P}}$, but with a more dramatic shift in the current-voltage curve for smaller $R_{\mathrm{P}}$.

The fitted values $k = 350\,\mathrm{pS}$ and $R_{\mathrm{P}} = 0.7\,\mathrm{nm}$ give reasonable agreement with the experimental data. Although the fitted intrinsic channel conductance $k$ may appear much larger than the estimate 170 pS reported in Ref. [12], $k$ represents the bare conductance before current self-responses are included. With the parameters used here, we obtain $k_{\mathrm{eff}} = 210\,\mathrm{pS}$ from Eq. (S.64), which is comparable to the estimate of 170 pS in Ref. [12].

Table S1 summarizes the fixed and derived parameter values used in the simulations. Although some parameters, such as the ionic diffusivity, depend on temperature, we keep them fixed because their variation

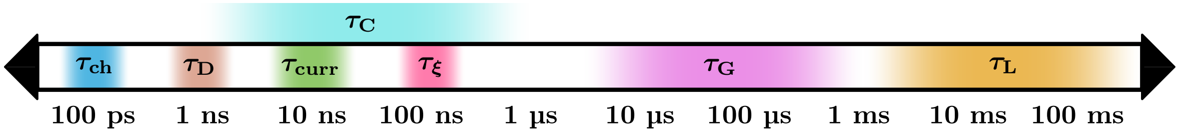

Figure S5: The range of timescales relevant to systems of single and multiple channels. These are the pore diffusion timescale $\tau_{\mathrm{ch}}$, the Debye timescale $\tau_{\mathrm{D}}$, the current adjustment timescale $\tau_{\mathrm{curr}}$, the gating drag timescale $\tau_{\xi}$, the capacitive timescale $\tau_{\mathrm{C}}$, the channel transition timescale $\tau_{\mathrm{G}}$, and the salt diffusion timescale $\tau_{\mathrm{L}}$. We neglect salt diffusion in our analysis and invoke the separation between $\tau_{\mathrm{G}}$ and all smaller timescales in the mean-field analysis of Sec. V.

is small over the temperature range considered here, $230\,\mathrm{K} < T < 320\,\mathrm{K}$.

##### 5. Cascade of timescales

In the preceding sections, we have identified a wide range of timescales that control the single-channel response. It may be useful to summarize the timescales relevant to this study. Arranged approximately from shortest to longest, these are

- **The pore diffusion timescale** $\tau_{\mathrm{ch}} \sim 0.1\,\mathrm{ns}$: The timescale for ionic diffusion over the width of the channel pore $R_{\mathrm{P}}$.
- **The Debye timescale** $\tau_{\mathrm{D}} \sim 1\,\mathrm{ns}$: The characteristic timescale for diffusion over the Debye length $\lambda_{\mathrm{D}}$ that controls reorganization within an electrical double layer.
- **The current adjustment timescale** $\tau_{\mathrm{curr}} \sim 10\,\mathrm{ns}$: The timescale over which the current through a single channel adjusts to a new steady state following a perturbation to the surrounding environment or internal channel structure.
- **The gating-variable drag timescale** $\tau_{\xi} \sim 100\,\mathrm{ns}$: The timescale required for the gating variable $\xi$ to sufficiently sample an energy well and approach an equilibrium distribution of states.
- **The capacitive timescale** $\tau_{\mathrm{C}} \sim 1$–$10^4$ ns: The timescale required to reach a new steady state following a perturbation of the applied potential, which is also the timescale required for a single-channel propagating electrical signal to reach the electrode separation distance $L$.
- **The bare channel transition timescale** $\tau_{\mathrm{G}} \sim 10$–$1000\,\mu\mathrm{s}$: The average timescale for a channel to move between the open and closed states when maintained at its gating voltage $V_{\mathrm{G}}$.
- **The salt diffusion timescale** $\tau_{\mathrm{L}} \sim 1$–$100\,\mathrm{ms}$: The timescale for salt diffusion to reach the electrode separation distance $L$, beyond which the slab geometry and electrode reactions affect the salt concentration response near the membrane.

Figure S5 summarizes graphically this spectrum of timescales. We note that the channel transition timescale $\tau_{\mathrm{G}}$ is longer than all other single-channel timescales, often by several orders of magnitude. This reflects the high energy barrier $\beta U^{\ddagger} \gg 1$ between the two energy wells corresponding to the open and closed states. This timescale separation allows the faster electrochemical response and local gating relaxation to be treated as effectively equilibrated when analyzing transitions between the open and closed states. It also provides the basis for the mean-field analysis in Sec. V.

#### II. Multiple channels

We now consider a membrane containing multiple identical channels of the same type. The channels interact indirectly through the electrochemical fields induced by their local transmembrane currents. Specifically, each channel modifies the membrane potential, charge density, and salt concentration experienced by the other channels, thereby affecting their gating and current. We retain the notation introduced for a single channel, with an additional subscript $i = 1, \ldots, N$ labeling the channels. At each channel, we track the gating variable $\xi_i$, the current $I_i$, the membrane potential $V_i$, the charge density $\rho_i$, and the salt concentration perturbation $C_i$.

##### 1. Gating dynamics

Each channel independently follows the same gating kinetics introduced for a single channel in Sec. I.1. The total gating energy can be written as

$$U(\xi_1, \ldots, \xi_N) \;=\; \sum_{i=1}^{N} \left[ U^{\ddagger} \left( \xi_i^4 - 2\xi_i^2 \right) - Q\,(V_i - V_{\mathrm{G}})\,\xi_i \right] \;. \tag{S.69}$$

The dynamics of the gating variables $\xi_i$ of the individual channels are governed by the overdamped Langevin equations:

$$\zeta\dot{\xi}_i \;=\; -\frac{\partial U}{\partial \xi_i} + \eta_i(t) \;=\; -4U^{\ddagger}\left(\xi_i^3 - \xi_i\right) + Q\left(V_i - V_{\mathrm{G}}\right) + \eta_i(t) \;, \tag{S.70}$$

where $\eta_i(t)$ is Gaussian white noise with $\langle \eta_i(t)\rangle = 0$ and $\langle \eta_i(t)\eta_j(t')\rangle = 2k_{\mathrm{B}}T\zeta\delta_{ij}\delta(t-t')$. These stochastic equations are coupled through the membrane potentials $V_i$, which depend on the current histories of all channels and therefore on their gating states.

#### 2. Electrochemical interaction between channels

The current through each channel is given by the same local current law as Eq. (S.56):

$$I_i(t) \;=\; -\Theta\left(\xi_i(t)\right) kV_i(t)\left[1 + \frac{1}{2C_0}\frac{C_i(t) + \rho_i(t)/\mathrm{e}}{\tanh\left(\beta_{\mathrm{ref}}\mathrm{e}V_i(t)/2\right)}\right] \;. \tag{S.71}$$

Although the current through channel $i$ is determined by the local quantities $V_i$, $\rho_i$, and $C_i$, these quantities contain nonlocal contributions arising from the currents through all channels. In the linear regime of the electrochemical response, these contributions simply superpose. Denoting the position of channel $i$ in the membrane by $\boldsymbol{x}_i$ and the distance between channels by $r_{ij} = |\boldsymbol{x}_i - \boldsymbol{x}_j|$, we then have

$$V_i(t) \;=\; \frac{\chi}{\chi+1}V_{\mathrm{ext}} + \sum_{j=1}^{N}\int_{-\infty}^{t} I_j(\tau)G_{\mathrm{V}}(r_{ij}, t-\tau)\,d\tau \;, \tag{S.72}$$

$$\rho_i(t) \;=\; \frac{g}{2(\chi+1)D}V_{\mathrm{ext}} + \sum_{j=1}^{N}\int_{-\infty}^{t} I_j(\tau)G_{\rho}(r_{ij}, t-\tau)\,d\tau \;, \tag{S.73}$$

$$C_i(t) \;=\; \sum_{j=1}^{N}\int_{-\infty}^{t} I_j(\tau)G_{\mathrm{C}}(r_{ij}, t-\tau)\,d\tau \;, \tag{S.74}$$

where the terms proportional to $V_{\mathrm{ext}}$ account for the applied electrode potential and the summations account for the electrochemical perturbations produced by all channel currents. In each summation, the $j = i$ contribution corresponds to the self-response from the single-channel model, while the $j \neq i$ contributions describe interactions between distinct channels. As noted above, these interactions are nonlocal and history-dependent, since the Green's functions extend over finite spatial and temporal ranges before decaying.

##### (a). Exponential approximation of electrochemical effects

As in the single-channel model, we again approximate the Green's functions by exponential functions to construct a numerically tractable framework for the multiple-channel dynamics. For the membrane potential and charge density, electrode screening imposes an effective cutoff distance $L$, so that $\tilde{G}_{\mathrm{V}}(r,t) = \tilde{G}_{\rho}(r,t) = 0$ for $r > L$. Within the cutoff distance, the electrical propagation time satisfies $\tau_{\mathrm{V}}(r) = \max(\tau_{\mathrm{D}}, r/v) \leq \tau_{\mathrm{V}}(L) = \tau_{\mathrm{C}}$. For the electrode separation distances $L \lesssim 5\,\mu\mathrm{m}$ considered here, the upper bound is $\tau_{\mathrm{C}} \sim 100\,\mathrm{ns}$. Even for larger values of $L$, such as in the squid giant axon, where the electrode separation is on the order of $L \sim 2\,\mathrm{mm}$ [32], the resulting capacitive timescale is estimated as $\tau_{\mathrm{C}} \sim 3\,\mu\mathrm{s}$ [30]. This timescale is shorter than the channel transition timescale $\tau_{\mathrm{G}} \sim 10$–$1000\,\mu\mathrm{s}$, which defines the scale of the mean residence time in a gating state. Therefore, when describing channel interactions that affect gating, the detailed propagation delay of the electrical response is not central to the dynamics. We therefore approximate $\tau_{\mathrm{V}}(r)$ by the local Debye relaxation time $\tau_{\mathrm{D}} \sim 1\,\mathrm{ns}$. Physically, this treatment means that the electrical perturbation mediating channel interactions is taken to reach any channel within the cutoff region on the Debye timescale $\tau_{\mathrm{D}}$, much faster than channel opening and closing.

Accordingly, Eqs. (S.72)–(S.74) become

$$V_i(t) = \underbrace{\frac{\chi}{\chi+1}V_{\text{ext}}}_{\text{applied}} + \underbrace{\frac{1}{\pi g\lambda_{\text{D}}}\int_{-\infty}^{t} I_i(\tau)\frac{e^{-(t-\tau)/\tau_{\text{D}}}}{\tau_{\text{D}}}\,d\tau}_{\text{self}} + \underbrace{\sum_{\substack{j=1\\0<r_{ij}<L}}^{N} \frac{\chi}{1+\chi}\frac{1}{\pi g r_{ij}}\int_{-\infty}^{t} I_j(\tau)\frac{e^{-(t-\tau)/\tau_{\text{D}}}}{\tau_{\text{D}}}\,d\tau}_{\text{distinct}} \ , \tag{S.75}$$

$$\rho_i(t) = \underbrace{\frac{g}{2(\chi+1)D}V_{\text{ext}}}_{\text{applied}} + \underbrace{\frac{1}{\pi D R_{\text{P}}}\int_{-\infty}^{t} I_i(\tau)\frac{e^{-(t-\tau)/\tau_{\text{ch}}}}{\tau_{\text{ch}}}\,d\tau}_{\text{self}} + \underbrace{\sum_{\substack{j=1\\0<r_{ij}<L}}^{N} \frac{1}{1+\chi}\frac{1}{2\pi D r_{ij}}\int_{-\infty}^{t} I_j(\tau)\frac{e^{-(t-\tau)/\tau_{\text{D}}}}{\tau_{\text{D}}}\,d\tau}_{\text{distinct}} \ , \tag{S.76}$$

$$C_i(t) = \underbrace{\frac{1}{\pi D R_{\text{P}}\text{e}}\int_{-\infty}^{t} I_i(\tau)\frac{e^{-(t-\tau)/\tau_{\text{ch}}}}{\tau_{\text{ch}}}\,d\tau}_{\text{self}} + \underbrace{\sum_{\substack{j=1\\j\neq i}}^{N} \frac{1}{2\pi D r_{ij}\text{e}}\int_{-\infty}^{t} I_j(\tau)\frac{e^{-(t-\tau)/\tau_{\text{diff}}(r_{ij})}}{\tau_{\text{diff}}(r_{ij})}\,d\tau}_{\text{distinct}} \ , \tag{S.77}$$

where $\tau_{\text{diff}}(r) = \pi^2 r^2/(4D)$ is the diffusion timescale over distance $r$, introduced in Sec. I.3. Here, the applied terms account for the externally imposed electrode potential, the self terms are the $j = i$ contributions corresponding to the self-response of each channel to its own current, and the distinct terms describe interactions between distinct channels.

#### (b). Scaling of the salt concentration response

While electrochemical interactions mediated by the membrane potential and charge density are restricted to channels within the cutoff distance $L$, the salt concentration perturbation spreads diffusively and therefore has no comparable finite cutoff scale. * Nevertheless, the following scaling argument shows that these contributions may be neglected over the simulation time window.

Let us consider an infinite membrane with channels separated by a lattice spacing $\ell$. To estimate the largest possible salt concentration perturbation generated by other channels, we let all channels open at $t = 0$ and carry the same constant current $I$. The nonself salt concentration perturbation at channel $i$ is then

$$\sum_{j\neq i}\int_0^t I\,G_{\text{C}}(r_{ij}, t-\tau)\,d\tau \;=\; I\sum_{j\neq i}\int_0^t G_{\text{C}}(r_{ij},\tau)\,d\tau \ , \tag{S.78}$$

which is independent of $i$ by translational invariance. Approximating the sum over channels by an integral over the membrane plane gives

$$\sum_{j\neq i}\int_0^t I\,G_{\text{C}}(r_{ij}, t-\tau)\,d\tau \;\approx\; \frac{I}{\ell^2}\int_0^t\int_{\Lambda}^{\infty} G_{\text{C}}(r,\tau)\,2\pi r\,dr\,d\tau \ . \tag{S.79}$$

Here, $\Lambda \sim \ell$ is a near-field cutoff that excludes the self-response. Since the integral remains finite as $\Lambda \to 0$, we set $\Lambda = 0$ to obtain an upper-bound estimate for the contribution from other channels. Using the salt concentration Green's function $G_{\text{C}}$ in Eq. (S.43), we obtain

$$\sum_{j\neq i}\int_0^t I\,G_{\text{C}}(r_{ij}, t-\tau)\,d\tau \;\approx\; \frac{I}{\ell^2}\int_0^t\int_0^{\infty} \frac{2}{\text{e}}\frac{e^{-r^2/(4D\tau)}}{(4\pi D\tau)^{3/2}}\,2\pi r\,dr\,d\tau \;=\; \frac{2I}{\ell^2\text{e}}\sqrt{\frac{t}{\pi D}} \ , \tag{S.80}$$

which grows as $\sqrt{t}$, characteristic of one-dimensional diffusion resulting from integration over the two-dimensional distribution of channels. The same result is obtained if the approximate kernel $\tilde{G}_{\text{C}}$ in Eq. (S.47) is used in place of $G_{\text{C}}$.

---

*In practice, including salt concentration perturbations generated by all other channels would therefore substantially increase the computational cost for large channel populations.

We now ask when the salt concentration perturbation from other channels in Eq. (S.80) becomes significant. Using the linearized current relation $I = -k_{\rm eff} V_{\rm ext}$ from Eq. (S.63), the concentration perturbation reaches the initial salt concentration $2C_0$ at $t \sim 4\tau_{\rm L}/(4\alpha \hat{V}_{\rm ext})^2$. Here, $\alpha \equiv (k_{\rm eff}/\ell^2)/(g/L)$ is the ratio of trans-membrane conductance to bulk conductance, and $\hat{V}_{\rm ext} = \beta_{\rm ref} e[\chi/(\chi+1)]V_{\rm ext}/2$ is the applied voltage scaled by the thermal voltage. The dimensionless parameter $\alpha$ represents the strength of current-induced interactions between channels: for a given bulk conductance $g/L$, larger $\alpha$ corresponds to a larger current through the membrane and therefore leads to stronger electrochemical perturbations. Thus, the salt concentration perturbations from other channels become significant on the salt diffusion timescale $\tau_{\rm L}$, with a larger applied voltage or stronger current-induced interactions further reducing this timescale. The two dimensionless parameters $\alpha$ and $\hat{V}_{\rm ext}$ naturally reappear in the mean-field analysis of Sec. V.

For realistic channel parameters, $\alpha, \hat{V}_{\rm ext} \sim 1$, so $\tau_{\rm L} \sim L^2/D$. With electrode separation distances $L \approx 1$–$10\,\mu$m and ionic diffusivity $D \approx 1\,{\rm nm}^2/{\rm ns}$, we obtain $\tau_{\rm L} \approx 1$–$100\,{\rm ms}$. As shown in Fig. S5, the salt diffusion timescale is much longer than the other timescales in the model. The collective behaviors underlying the phase transitions in the finite systems simulated here occur on timescales much shorter than this slow salt-diffusion timescale. Since the simulations are performed over times that do not exceed this scale, we neglect salt concentration perturbations generated by other channels. Beyond $t \gtrsim \tau_{\rm L}$, large salt concentration differences may build up across the membrane, producing significant osmotic pressure. Such pressure could rupture reconstituted membranes [33] or activate pressure-relief mechanisms in real cells [34], neither of which is included in the present model.

##### (c). History variable formulation

With the simplification applied to the concentration effects, we now write Eqs. (S.75)–(S.77) in terms of quantities common to all three equations. In each summation for distinct channel interactions, the time integral depends only on the current through the source channel $j$ and not on the target channel $i$. Therefore, we introduce two "history variables" for each channel, which encode the memory of the current:

$$H_i(t) = \int_{-\infty}^{t} I_i(\tau) \frac{e^{-(t-\tau)/\tau_{\rm D}}}{\tau_{\rm D}}\, d\tau \,, \qquad F_i(t) = \int_{-\infty}^{t} I_i(\tau) \frac{e^{-(t-\tau)/\tau_{\rm ch}}}{\tau_{\rm ch}}\, d\tau \,. \tag{S.81}$$

Here, the current history $I_i(t)$ is convolved with exponential memory kernels, $e^{-t/\tau_{\rm D}}/\tau_{\rm D}$ for $H_i$ and $e^{-t/\tau_{\rm ch}}/\tau_{\rm ch}$ for $F_i$. Current history older than the corresponding relaxation timescale, $\tau_{\rm D}$ or $\tau_{\rm ch}$, is exponentially suppressed, so earlier currents have a weaker influence on the present interaction. For a constant current $I$ sustained over a time much longer than these timescales, the transient part of the current history is suppressed, leaving only the steady current: $H_i, F_i \to I$.

Substituting the history variables into Eqs. (S.75)–(S.77), we obtain

$$V_i(t) = \frac{\chi}{\chi+1} V_{\rm ext} + \frac{H_i(t)}{\pi g \lambda_{\rm D}} + \sum_{\substack{j=1 \\ 0<r_{ij}<L}}^{N} \frac{\chi}{1+\chi} \frac{H_j(t)}{\pi g r_{ij}} \,, \tag{S.82}$$

$$\rho_i(t) = \frac{g}{2(\chi+1)D} V_{\rm ext} + \frac{F_i(t)}{\pi D R_{\rm P}} + \sum_{\substack{j=1 \\ 0<r_{ij}<L}}^{N} \frac{1}{1+\chi} \frac{H_j(t)}{2\pi D r_{ij}} \,, \tag{S.83}$$

$$C_i(t) = \frac{F_i(t)}{\pi D R_{\rm P} e} \,, \tag{S.84}$$

where the distinct-channel salt concentration contributions to $C_i$ have been neglected based on the scaling argument in the previous section. In this reduced formulation, interactions between distinct channels enter only through the electrochemical response encoded in $H_j(t)$, which occurs on the Debye timescale $\tau_{\rm D}$.

#### 3. Summary of the multi-channel model

We now present the equations used for the multiple-channel simulations and summarize the assumptions underlying the model. For each channel, the model evolves the gating variable $\xi_i$, current $I_i$, membrane potential $V_i$, charge density $\rho_i$, salt concentration perturbation $C_i$, and history variables $H_i$ and $F_i$. Combining the gating dynamics, current law, electrochemical response, and history-variable formulation gives the following reduced model:

$$\zeta \frac{d\xi_i(t)}{dt} = -4U^{\ddagger}\left(\xi_i(t)^3 - \xi_i(t)\right) + Q\left(V_i(t) - V_{\rm G}\right) + \eta_i(t) \ , \tag{S.85}$$

$$\langle \eta_i(t) \rangle = 0 \ , \qquad \left\langle \eta_i(t)\eta_j(t') \right\rangle = 2\zeta k_{\rm B} T \delta(t-t')\delta_{ij} \ , \tag{S.86}$$

$$I_i(t) = -\Theta\left(\xi_i(t)\right) k V_i(t) \left[1 + \frac{1}{2C_0} \frac{C_i(t) + \rho_i(t)/{\rm e}}{\tanh\left(\beta_{\rm ref} {\rm e} V_i(t)/2\right)}\right] \ , \tag{S.87}$$

$$V_i(t) = \frac{\chi}{\chi+1} V_{\rm ext} + \frac{H_i(t)}{\pi g \lambda_{\rm D}} + \sum_{\substack{j=1 \\ 0<r_{ij}<L}}^{N} \frac{\chi}{1+\chi} \frac{H_j(t)}{\pi g r_{ij}} \ , \tag{S.88}$$

$$\rho_i(t) = \frac{g}{2(\chi+1)D} V_{\rm ext} + \frac{F_i(t)}{\pi D R_{\rm P}} + \sum_{\substack{j=1 \\ 0<r_{ij}<L}}^{N} \frac{1}{1+\chi} \frac{H_j(t)}{2\pi D r_{ij}} \ , \tag{S.89}$$

$$C_i(t) = \frac{F_i(t)}{\pi D R_{\rm P} {\rm e}} \ , \tag{S.90}$$

$$H_i(t) = \int_{-\infty}^{t} I_i(\tau) \frac{e^{-(t-\tau)/\tau_{\rm D}}}{\tau_{\rm D}} d\tau \ , \tag{S.91}$$

$$F_i(t) = \int_{-\infty}^{t} I_i(\tau) \frac{e^{-(t-\tau)/\tau_{\rm ch}}}{\tau_{\rm ch}} d\tau \ . \tag{S.92}$$

In the simulations, we use the parameters listed in Table S1 and introduce three additional multiple-channel parameters: the electrode separation $L$, which determines the interaction cutoff, the channel spacing $\ell$, and the number of channels $N$.

The reduced model is constructed under the following assumptions. Each assumption may be relaxed in future extensions of this work:

- *Gating*:
  - Channel gating is described by a two-state model, with open and closed states separated by a single energy barrier.
  - Changes in membrane potential modify the gating energy linearly, resulting in a sigmoidal open probability.
  - Channel gating depends only on the local membrane potential.
- *Current-voltage law*:
  - Ions within a channel are sufficiently dilute to be treated as non-interacting.
  - The number of barriers in a channel is sufficiently large that the approximation in Eq. (S.21) can be used.
  - Current adjustment is fast compared with channel gating, so the current is treated as quasi-steady and its value is determined by the local electrochemical environment.
- *Electrochemical response*:
  - Each Green's function is well approximated by a single exponential function with the appropriate magnitude and timescale.

- Membrane potential and charge density perturbations produced by a channel are negligible beyond the electrode separation distance $L$.
- Membrane-potential and charge-density signals between interacting channels propagate on the timescale $\tau_{\mathrm{D}}$.
- Salt concentration perturbations from other channels are negligible over the simulation time window.

### III. Numerical implementation

We provide details of the numerical implementation used to simulate the system of equations given by Eqs. (S.85)–(S.92). This includes the time-stepping methods used for the stochastic differential equation and the history-variable integrals.

**Stochastic ODE:** To evolve the stochastic ODE governing $\xi_i$, the Euler–Maruyama method is used [35]. Integrating from time $t$ to $t + \Delta t$ gives

$$\zeta\left[\xi_i(t+\Delta t) - \xi_i(t)\right] \;\approx\; -\Delta t\left[4U^{\ddagger}(\xi_i(t)^3 - \xi_i(t)) - Q(V_i(t) - V_{\mathrm{G}})\right] + \int_t^{t+\Delta t} d\tau\, \eta_i(\tau)\ . \tag{S.93}$$

The last term is sampled from the statistics of $\eta_i$ specified in Eq. (S.86), giving

$$\int_t^{t+\Delta t} d\tau\, \eta_i(\tau) \;\sim\; \mathcal{N}(0, 2k_{\mathrm{B}}T\zeta\Delta t)\ , \tag{S.94}$$

where $\mathcal{N}$ denotes the normal distribution, here with mean 0 and variance $2k_{\mathrm{B}}T\zeta\Delta t$.

**History variables:** To integrate the history variables from time $t$ to $t + \Delta t$, we rely on the semigroup property of the exponential function to find

$$\begin{aligned} H_i(t+\Delta t) \;&=\; \int_{-\infty}^{t+\Delta t} d\tau\, I_i(\tau)\frac{e^{-(t+\Delta t-\tau)/\tau_{\mathrm{D}}}}{\tau_{\mathrm{D}}} \\ &=\; e^{-\Delta t/\tau_{\mathrm{D}}}\int_{-\infty}^{t} d\tau\, I_i(\tau)\frac{e^{-(t-\tau)/\tau_{\mathrm{D}}}}{\tau_{\mathrm{D}}} + \int_t^{t+\Delta t} d\tau\, I_i(\tau)\frac{e^{-(t+\Delta t-\tau)/\tau_{\mathrm{D}}}}{\tau_{\mathrm{D}}}\ , \end{aligned} \tag{S.95}$$

where we recognize that the first integral is precisely $H_i(t)$. Therefore, we have the exact relation

$$H_i(t+\Delta t) \;=\; e^{-\Delta t/\tau_{\mathrm{D}}} H_i(t) + \int_t^{t+\Delta t} d\tau\, I_i(\tau)\frac{e^{-(t+\Delta t-\tau)/\tau_{\mathrm{D}}}}{\tau_{\mathrm{D}}}\ , \tag{S.96}$$

where $H_i(t + \Delta t)$ no longer depends explicitly on the entire history of $I_i(t)$, but only on its values over the previous time step. Such a formulation is only possible because we approximated the memory kernel as exponential in time.* The original functions, which include noninteger powers, are not amenable to a recurrence. This strategy is analogous to using sums of exponential kernels (Prony series) to represent viscoelastic moduli [36].

We approximate the integral from $t$ to $t + \Delta t$ using a forward approximation, with $I_i(\tau) \approx I_i(t)$ over the interval. This yields

$$H_i(t+\Delta t) \;\approx\; e^{-\Delta t/\tau_{\mathrm{D}}} H_i(t) + \left(1 - e^{-\Delta t/\tau_{\mathrm{D}}}\right) I_i(t)\ , \tag{S.97}$$

where the time-stepping procedure can be interpreted as a weighted average of the previous value of the history variable and the current value of the current. Applying the same approach to $F_i$ yields an identical recurrence with $\tau_{\mathrm{ch}}$ instead of $\tau_{\mathrm{D}}$.

*Similar recurrences are only possible when the kernel is exponential (as it is here), sinusoidal, polynomial, or an algebraic combination of such functions.

**Governing equations:** We discretize time into uniform steps of $\Delta t$, with $K$ total time steps. We use the superscript $\alpha = 0, 1, \ldots, K$ to denote the values of quantities at time $t^\alpha = \alpha\Delta t$. Therefore, the scalar quantities we solve for numerically are $\xi_i^\alpha$, $I_i^\alpha$, $V_i^\alpha$, $\rho_i^\alpha$, $C_i^\alpha$, $H_i^\alpha$, and $F_i^\alpha$ for $1 \le i \le N$ and $0 \le \alpha \le K$, with initial conditions for $\xi_i^0$, $H_i^0$, and $F_i^0$.

We divide the quantities into two groups. First, there are those that are *quasi-static*, i.e., determined only by the present values of the other quantities. These are the membrane potential, the charge density, the salt concentration, and the current. Second, there are the *time-evolving* quantities, which have dynamical governing equations. These are the gating variable and the history variables. At each time step, including $t = 0$, we compute the quasi-static quantities as follows:

$$I_i^\alpha = -\Theta(\xi_i^\alpha) k V_i^\alpha \left[1 + \frac{1}{2C_0} \frac{C_i^\alpha + \rho_i^\alpha/\mathrm{e}}{\tanh\!\big(\beta_{\mathrm{ref}} \mathrm{e} V_i^\alpha/2\big)}\right] , \tag{S.98}$$

$$V_i^\alpha = \frac{\chi}{\chi+1} V_{\mathrm{ext}} + \frac{H_i^\alpha}{\pi g \lambda_{\mathrm{D}}} + \sum_{\substack{j=1 \\ 0<r_{ij}<L}}^{N} \frac{\chi}{1+\chi} \frac{H_j^\alpha}{\pi g r_{ij}} , \tag{S.99}$$

$$\rho_i^\alpha = \frac{g}{2(\chi+1)D} V_{\mathrm{ext}} + \frac{F_i^\alpha}{\pi D R_{\mathrm{P}}} + \sum_{\substack{j=1 \\ 0<r_{ij}<L}}^{N} \frac{1}{1+\chi} \frac{H_j^\alpha}{2\pi D r_{ij}} , \tag{S.100}$$

$$C_i^\alpha = \frac{F_i^\alpha}{\pi D R_{\mathrm{P}} \mathrm{e}} , \tag{S.101}$$

where we first compute $V_i^\alpha$, $\rho_i^\alpha$, and $C_i^\alpha$ before computing $I_i^\alpha$. Here, we assume that the time-evolving quantities, $\xi_i^\alpha$, $H_i^\alpha$, and $F_i^\alpha$, have already been computed at $t^\alpha$ from the previous time integration step (or are prescribed as the initial conditions for $\alpha = 0$). Next, we proceed to the time integration step for each $\alpha \ge 0$, given by

$$\xi_i^{\alpha+1} = \xi_i^\alpha - \frac{\Delta t}{\zeta}\left[4U^\ddagger \left((\xi_i^\alpha)^3 - \xi_i^\alpha\right) - Q(V_i^\alpha - V_{\mathrm{G}})\right] + \sqrt{\frac{2k_{\mathrm{B}}T\Delta t}{\zeta}}\,\eta_i^\alpha , \qquad \eta_i^\alpha \sim \mathcal{N}(0,1) , \tag{S.102}$$

$$H_i^{\alpha+1} = e^{-\Delta t/\tau_{\mathrm{D}}} H_i^\alpha + \left(1 - e^{-\Delta t/\tau_{\mathrm{D}}}\right) I_i^\alpha , \tag{S.103}$$

$$F_i^{\alpha+1} = e^{-\Delta t/\tau_{\mathrm{ch}}} F_i^\alpha + \left(1 - e^{-\Delta t/\tau_{\mathrm{ch}}}\right) I_i^\alpha , \tag{S.104}$$

where each $\eta_i^\alpha$ is sampled independently. In the time integration step, each of the three time-evolving quantities at step $\alpha + 1$ is computed from values at the previous time step $\alpha$.

Overall, the time-stepping framework consists of computing the quasi-static quantities from the time-evolving ones at each time point, and then, at each time step, integrating the time-evolving quantities to new values. Of particular importance is that the quasi-static quantities involve the interactions between channels; for example, in the governing equations for $V_i^\alpha$ (Eq. (S.99)) and $\rho_i^\alpha$ (Eq. (S.100)), the values of the history variables at other channels $j$ are used. However, during the time integration step, all operations are local, with the integration of the gating and history variables constrained exclusively to quantities at channel $i$.

**OpenMM implementation:** The molecular dynamics package `OpenMM` is used to implement the above time-stepping scheme. Within the `OpenMM` framework, each channel is considered a "particle" on the two-dimensional surface of the membrane. `OpenMM` provides support for particle positions, velocities, and forces, but not for additional per-particle quantities. Accordingly, we have developed an extension of `OpenMM` with *generalized coordinates*, as in simulations of dynamical systems, which allows any number of additional quantities for each particle—in this case, seven per particle. These generalized coordinates are accessible both in force calculations and during time integration, analogous to the need for positions and velocities in these steps.

Next, although this system does not have traditional forces or molecular dynamics, we associate each quantity with either "force" calculations or "time integration" calculations. The natural subdivisions are exactly the quasi-static and time-evolving groups we identified in Sec. III. As an analogy, forces in a molecular system are determined by the instantaneous conditions in a quasi-static manner and couple all particles

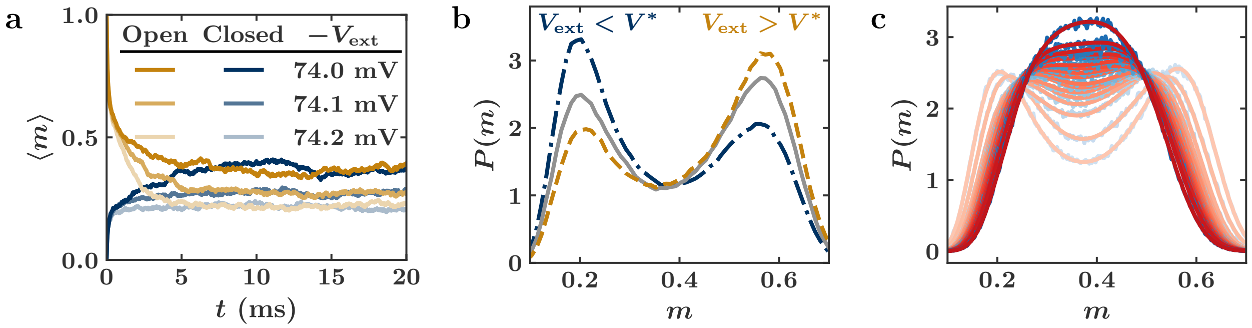

Figure S6: (a) The convergence of the replica-averaged open fraction $\langle m \rangle$ in time, from initially open (gold) and closed (blue) states. Simulations are shown for $N = 20^2$ and $T = 270\,\mathrm{K}$. (b) The distributions of the open fraction over equilibrated samples for $N = 20^2$ and $T = 270\,\mathrm{K}$ at applied potentials of $-74.0\,\mathrm{mV}$ (blue), $-73.97\,\mathrm{mV}$ (gray), and $-73.94\,\mathrm{mV}$ (gold). Bimodality indicates two distinct phases (mostly open and mostly closed) separated by a first-order phase transition, with the middle value of $V_{ext}$ identified as the coexistence potential $V^*(T)$. (c) Histograms of the open fraction at the identified coexistence potential from $T = 270\,\mathrm{K}$ (lightest curves) to $T = 290\,\mathrm{K}$ (darkest curves). The transition from two peaks to one peak indicates the existence of a critical point where the distribution flattens. Blue curves show the raw histogram data obtained from sampling, and red curves show the results obtained by fitting a free energy.

together, just as the membrane potential and charge density are determined by the history variables at the other channels at a given time. Meanwhile, in a molecular system, the evolution of a particle's position and velocity involves time integration and is based only on the forces present at the particle, just as the gating and history variables evolve based only on local quantities at the given channel.

With this association in mind, we have created a plugin for `OpenMM` that implements custom `Force` and `Integrator` classes to perform the two halves of the time-stepping scheme described above. The `IonChannelForce` class computes the quasi-static quantities from the time-evolving ones at any time. The `IonChannelIntegrator` class advances the time-evolving quantities from time $t$ to time $t + \Delta t$. While the model does not describe conservative forces typical in molecular dynamics simulations, the workflow for the multi-channel problem is analogous in structure, which is why we choose to use `OpenMM` to implement it.

### IV. Data analysis

We provide details of the calculations used to obtain statistics for the analysis of the first-order phase transition shown in Fig. 2 of the main text and for the analysis of the critical point shown in Fig. 3 of the main text.

#### 1. First-order transition analysis

All simulations are performed with a time step of $\Delta t = 0.1\,\mathrm{ns}$ and are run to a final time of at least $20\,\mathrm{ms}$. For all simulation conditions, half of the replicates are initialized in the all-open state and half in the all-closed state. Open fractions reported in the manuscript are both replica- and time-averaged starting from the estimated convergence time. Figure S6a shows the convergence of the open fraction $m$ over $20\,\mathrm{ms}$ in a $20 \times 20$ system at $T = 270\,\mathrm{K}$ for three values of $V_{ext}$.

Susceptibility is computed using the fluctuation formula

$$\chi = N\beta \left( \langle m^2 \rangle - \langle m \rangle^2 \right) , \tag{S.105}$$

where $m$ is the open fraction of channels and $\langle \cdot \rangle$ denotes both time and replica averaging, starting after the convergence time.

Probability density functions for the open fraction $m$ in a finite system are calculated as histograms over the $N + 1$ possible values of the open fraction for a system with $N$ channels. These curves are then further binned into approximately 100 equal-size groups for each system, depending on the value of $N$, to reduce the fluctuations that arise from sampling each possible value separately. Figure S6b shows three histograms for a $20 \times 20$ system at three values of $V_{\text{ext}}$, where the curves are smoothed by binning the distribution of states into groups of 4 (e.g., 0–3 channels open, 4–7 open, etc.).

##### 2. Critical point analysis

The average open fraction $m$ and the probability density $P(m)$ are computed exactly as in the first-order phase transition analysis.

Using the histograms, a temperature $T$ is identified as being below the critical temperature $T_{\text{C}}$ if there exists a regime of $V_{\text{ext}}$ where the probability density is bimodal. For such temperatures, the coexistence condition is defined by the value of the applied potential $V_{\text{ext}}$ such that the two peaks have equal heights, denoted $V^*(T)$. In Fig. S6b, this corresponds to the gray curve, which is closest to having equal peak heights for $T = 270\,\text{K}$ among the three neighboring values of $V_{\text{ext}}$ plotted. The sensitivity of the estimate of the coexistence potential $V^*$ is $0.01\,\text{mV}$. The critical point is identified by the temperature at which the histogram loses its two-peak structure and becomes a single broad peak.

The values of the open fraction $m$ at the two peaks are found by fitting the histogram to an effective free energy obtained by assuming $f(m) = -\log(P(m))$. We use the following form for the free energy $f$:

$$f \;=\; -b(m\ln(m) + (1-m)\ln(1-m)) - \sum_{i=0}^{6} a_i m^i \;, \tag{S.106}$$

where the data are fitted for the eight parameters $b, a_0, \ldots, a_6$. The first term is included to capture the logarithmic behavior expected from entropy, while the others are a polynomial expansion in $m$ up to order 6. Order 6 is chosen since including terms only up to order 4 is consistent only with a critical point that is symmetric in $m$, whereas this system demonstrates an asymmetry. The free energy is constrained such that the histogram is normalized through $a_0$. Once this model is fitted, the two coexistence values of $m$ are found from the minima of $f$. Note that we fit the free energy since the noise in the histogram is too large to precisely find the peak locations in the full histogram.

Figure S6c shows the raw, unsmoothed histograms in blue from 270 K to 290 K at the identified coexistence potentials. The red curves show the fitted distributions using the above free energy. Extrema are extracted from the fitted free energy curves as described above. We observe very good agreement at every temperature between the numerically sampled histograms and the free energies fitted to these data.

#### V. Mean-field analysis

We perform a mean-field analysis to characterize the emergent phase-transition behavior of the multiple-channel system, whose dynamics are described in Sec. II.3. This analysis also identifies the key dimensionless groups that control the critical temperature and critical applied voltage in the $V_{\text{ext}}$–$T$ phase diagram. It also guides the choice of parameters when performing stochastic simulations to search for coexistence conditions between the collective open and closed states, as well as the critical point.

##### 1. Separation of timescales

We aim to analyze the steady-state behavior of the multiple-channel system, including the fraction of open channels and the corresponding current and membrane potential. To begin, note that the relaxation of the gating states occurs over the channel transition timescale $\tau_{\text{G}} \gtrsim 1\,\mu\text{s}$ (Fig. S5). In contrast, the history variables $H_i(t)$ and $F_i(t)$ in Eq. (S.81) encode how current history contributes to the present electrochemical response over the much shorter memory timescales $\tau_{\text{D}} \sim 1\,\text{ns}$ and $\tau_{\text{ch}} \lesssim 1\,\text{ns}$, respectively. This separation of timescales allows us to treat the electrochemical response as effectively instantaneous relative to changes in the transmembrane current on the timescale over which the gating states relax: $H_i(t), F_i(t) \approx I_i(t)$. Accordingly, the membrane potential, charge density, and salt concentration perturbation in Eqs. (S.88)–(S.90) become

$$V_i(t) \;=\; \frac{\chi}{\chi+1}V_{\rm ext} + \frac{I_i(t)}{\pi g\lambda_{\rm D}} + \sum_{\substack{j=1\\0<r_{ij}<L}}^{N}\frac{\chi}{1+\chi}\frac{I_j(t)}{\pi g r_{ij}}\;, \tag{S.107}$$

$$\rho_i(t) \;=\; \frac{g}{2(\chi+1)D}V_{\rm ext} + \frac{I_i(t)}{\pi D R_{\rm P}} + \sum_{\substack{j=1\\0<r_{ij}<L}}^{N}\frac{1}{1+\chi}\frac{I_j(t)}{2\pi D r_{ij}}\;, \tag{S.108}$$

$$C_i(t) \;=\; \frac{I_i(t)}{\pi D R_{\rm Pe}}\;, \tag{S.109}$$

The current constitutive law remains unchanged from Eq. (S.87):

$$I_i(t) \;=\; -n_i(t)kV_i(t)\left[1+\frac{1}{2C_0}\frac{C_i(t)+\rho_i(t)/{\rm e}}{\tanh\left(\beta_{\rm ref}{\rm e}V_i(t)/2\right)}\right]\;, \tag{S.110}$$

where we introduce $n_i \equiv \Theta(\xi_i(t))$ to indicate the channel state: open with $n_i = 1$ and closed with $n_i = 0$. For a given set of channel states $\{n_i\}$, the above set of $4N$ equations (S.107)–(S.110) constitutes a nonlinear system for $V_i$, $\rho_i$, $C_i$, and $I_i$. The channel-state configuration $\{n_i\}$ determines the local electrochemical variables, but only implicitly through the nonlinear system.

#### 2. Current through channels

We now apply a mean-field approximation to relate the current through an open channel to the fraction of open channels. This approximation amounts to replacing the electrochemical effects of the other channels by an average field that is generated by their mean current. Specifically, for a reference channel $i$, it allows us to replace the current through each of the other channels by a mean-field value $I_j = mI^{\rm o}_{\rm MF}$ in Eqs. (S.107)–(S.109), where $m$ is the fraction of open channels and $I^{\rm o}_{\rm MF}$ is the current through an open channel in the mean-field environment. We may then substitute the resulting mean-field expressions for $V_i$, $\rho_i$, and $C_i$ into the current law in Eq. (S.110). For self-consistency, however, the current law requires the reference channel to carry the mean-field open current, $I_i = I^{\rm o}_{\rm MF}$, when in the open state $n_i = 1$. This yields

$$I^{\rm o}_{\rm MF} \;=\; -k\left(\frac{\chi}{\chi+1}V_{\rm ext}+V_{\rm MF}+\frac{I^{\rm o}_{\rm MF}}{\pi g\lambda_{\rm D}}\right)\left[1+\frac{\frac{\beta_{\rm ref}{\rm e}}{2}\left(\frac{V_{\rm ext}}{\chi+1}+\frac{V_{\rm MF}}{\chi}+\frac{4I^{\rm o}_{\rm MF}}{\pi g R_{\rm P}}\right)}{\tanh\left(\frac{\beta_{\rm ref}{\rm e}}{2}\left(\frac{\chi}{\chi+1}V_{\rm ext}+V_{\rm MF}+\frac{I^{\rm o}_{\rm MF}}{\pi g\lambda_{\rm D}}\right)\right)}\right]\;, \tag{S.111}$$

where

$$V_{\rm MF} \;=\; \sum_{\substack{j=1\\0<r_{ij}<L}}^{N}\frac{\chi}{1+\chi}\frac{mI^{\rm o}_{\rm MF}}{\pi g r_{ij}}\;, \tag{S.112}$$

is the mean-field shift in the membrane potential induced by the other channels, and we have used the relation $\beta_{\rm ref} = g/(2C_0 D{\rm e}^2)$. The mean-field approximation decouples the interacting channels by incorporating the distinct-channel contribution into an effective imposed potential. Thus, Eq. (S.111) is indeed the single-channel current relation in Eq. (S.60), with the externally imposed potential replaced by $[\chi/(\chi+1)]V_{\rm ext} + V_{\rm MF}$.

We next nondimensionalize Eq. (S.111) to identify the governing dimensionless parameters. To that end, we first estimate the scale of the mean-field shift in Eq. (S.112) by treating the channels as distributed with a uniform density $\rho = \ell^{-2}$:

$$V_{\rm MF} \;\approx\; \frac{\chi}{1+\chi}\frac{mI^{\rm o}_{\rm MF}}{\pi g}\frac{1}{\ell^2}\int_0^L\frac{1}{r}\,2\pi r\,dr \;=\; \frac{\chi}{1+\chi}\frac{2L}{g\ell^2}mI^{\rm o}_{\rm MF}\;. \tag{S.113}$$

We then scale the open-channel current by the current that would flow through a channel under the external potential in the absence of any current-induced responses, $\tilde{I}^{\rm o}_{\rm MF} \equiv I^{\rm o}_{\rm MF}/(k[\chi/(\chi+1)]V_{\rm ext})$, and we scale the potentials by the reference thermal voltage, $\hat{V} \equiv \beta_{\rm ref}{\rm e}[\chi/(\chi+1)]V/2$. Accordingly, the current law in Eq. (S.111) can be nondimensionalized as

$$\tilde{I}^{\rm o}_{\rm MF} \;=\; -\left(1+2\tilde{I}^{\rm o}_{\rm MF}\left(\tilde{\gamma}+\tilde{\alpha}m\right)\right)\left[1+\frac{1}{\chi}\frac{\hat{V}_{\rm ext}\left(1+2\tilde{I}^{\rm o}_{\rm MF}\left(4\chi\frac{\lambda_{\rm D}}{R_{\rm P}}\tilde{\gamma}+\tilde{\alpha}m\right)\right)}{\tanh\left(\hat{V}_{\rm ext}\left(1+2\tilde{I}^{\rm o}_{\rm MF}\left(\tilde{\gamma}+\tilde{\alpha}m\right)\right)\right)}\right] \;, \tag{S.114}$$

where we define two dimensionless parameters,

$$\tilde{\alpha} \;=\; \frac{kV_{\rm MF}/2}{mI^{\rm o}_{\rm MF}} \;=\; \frac{\chi+1}{\chi}\frac{\hat{V}_{\rm MF}/\hat{V}_{\rm ext}}{2m\tilde{I}^{\rm o}_{\rm MF}} \;\approx\; \frac{\chi}{1+\chi}\frac{kL}{g\ell^2}\;, \qquad\qquad \tilde{\gamma} \;=\; \frac{k}{2\pi g\lambda_{\rm D}}\;. \tag{S.115}$$

The dimensionless parameter $\tilde{\alpha}$ characterizes the strength of interactions between channels, as introduced in Sec. II.2 (b). Here, it consistently quantifies the mean-field membrane potential shift relative to the mean current through the other channels. The other parameter $\tilde{\gamma}$ was already introduced in Sec. I.4, and it characterizes the strength of the self-response in the local membrane potential for a given externally imposed potential.

Equation (S.114) constitutes an implicit relation between the open-channel current $\tilde{I}^{\rm o}_{\rm MF}$ and the open fraction $m$, which generally requires numerical solution. Nevertheless, several properties of $\tilde{I}^{\rm o}_{\rm MF}$ are noteworthy. First, the mean-field current develops in the direction set by the externally imposed potential regardless of the current-induced corrections, i.e. $\tilde{I}^{\rm o}_{\rm MF}<0$. This follows by contradiction: if $\tilde{I}^{\rm o}_{\rm MF}$ were positive, the left-hand side of Eq. (S.114) would be positive, whereas the right-hand side would be negative.

Accordingly, self-consistency requires that the current-induced perturbations reduce the magnitude of the local membrane potential relative to the externally imposed potential, while preserving its sign. Thus, the factor $1+2\tilde{I}^{\rm o}_{\rm MF}(\tilde{\gamma}+\tilde{\alpha}m)$ in Eq. (S.114), which represents this relative local membrane potential, remains between 0 and 1. Together, we find bounds on the open-channel current, $-1/(2(\tilde{\gamma}+\tilde{\alpha}m))<\tilde{I}^{\rm o}_{\rm MF}<0$. In what follows, we examine two limiting regimes that admit explicit relations between $\tilde{I}^{\rm o}_{\rm MF}$ and $m$: small applied potentials, $|\hat{V}_{\rm ext}|\ll 1$, and large applied potentials, $|\hat{V}_{\rm ext}|\gg 1$.

#### (a). Small applied potentials

We first consider the regime $|\hat{V}_{\rm ext}|\ll 1$, where Eq. (S.114) admits a linearized form. Since the factor $1+2\tilde{I}^{\rm o}_{\rm MF}(\tilde{\gamma}+\tilde{\alpha}m)\in(0,1)$, as discussed above, the entire argument of the hyperbolic tangent remains small when $|\hat{V}_{\rm ext}|\ll 1$. Equation (S.114) then reduces to

$$\tilde{I}^{\rm o}_{\rm MF} \;=\; -\left(1+2\tilde{I}^{\rm o}_{\rm MF}\left(\tilde{\gamma}+\tilde{\alpha}m\right)\right)-\frac{1}{\chi}\left(1+2\tilde{I}^{\rm o}_{\rm MF}\left(4\chi\frac{\lambda_{\rm D}}{R_{\rm P}}\tilde{\gamma}+\tilde{\alpha}m\right)\right)\;. \tag{S.116}$$

Solving this linear equation for $\tilde{I}^{\rm o}_{\rm MF}$ yields an explicit relation between the mean-field current and the open fraction:

$$\tilde{I}^{\rm o}_{\rm MF} \;=\; -\frac{\chi+1}{\chi}\frac{\kappa}{1+2\alpha m}\;, \tag{S.117}$$

where

$$\kappa \;=\; \frac{k_{\rm eff}}{k} \;=\; \frac{1}{1+2\tilde{\gamma}\left(4\frac{\lambda_{\rm D}}{R_{\rm P}}+1\right)}\;, \qquad\qquad \alpha \;=\; \frac{\chi+1}{\chi}\kappa\tilde{\alpha} \;=\; \frac{k_{\rm eff}L}{g\ell^2}\;. \tag{S.118}$$

Here, $k_{\rm eff}=[1/k+1/(\pi g\lambda_{\rm D})+4/(\pi gR_{\rm P})]^{-1}$ is the effective channel conductance defined in Eq. (S.64), incorporating the apparent reduction of the channel conductance due to the electrochemical self-response in the linear regime.

This reduction also rescales the interaction parameter from $\tilde{\alpha}$ to $\alpha$ in the linearized relation. We note that while both $\tilde{\alpha}$ and $\alpha$ characterize the strength of collective interactions between channels, they connect different quantities. The parameter $\tilde{\alpha}$ relates the mean-field membrane potential shift $V_{\rm MF}$ to the mean current $mI^{\rm o}_{\rm MF}$. By contrast, once the current associated with this mean-field shift is evaluated using the effective conductance $k_{\rm eff}$, $\alpha$ directly governs the relation between the mean-field current $I^{\rm o}_{\rm MF}$ and the open fraction $m$. Consequently, $\alpha$ enters Eq. (S.117) through the denominator $1+2\alpha m$, which determines how strongly collective channel–channel interactions reduce the mean-field current as the open fraction $m$ increases. In dimensional form, Eq. (S.117) is given by

$$I^{\rm o}_{\rm MF} \;=\; -k_{\rm eff}\frac{V_{\rm ext}}{1+2\alpha m} \;. \tag{S.119}$$

##### (b). Large applied potentials

When the applied potential is large, $|\hat{V}_{\rm ext}| \gg 1$, the hyperbolic tangent term in Eq. (S.114) approaches $\mathrm{sgn}(\hat{V}_{\rm ext})$, leading to

$$\tilde{I}^{\rm o}_{\rm MF} \;=\; -\left(1+2\tilde{I}^{\rm o}_{\rm MF}\,(\tilde{\gamma}+\tilde{\alpha}m)\right)\left[1+\frac{|\hat{V}_{\rm ext}|}{\chi}\left(1+2\tilde{I}^{\rm o}_{\rm MF}\left(4\chi\frac{\lambda_{\rm D}}{R_{\rm P}}\tilde{\gamma}+\tilde{\alpha}m\right)\right)\right] \;, \tag{S.120}$$

which is a quadratic equation in $\tilde{I}^{\rm o}_{\rm MF}$. A dominant balance shows that $\tilde{I}^{\rm o}_{\rm MF} \sim \mathcal{O}(1)$ as $|\hat{V}_{\rm ext}| \to \infty$. This implies that the coefficient multiplying $|\hat{V}_{\rm ext}|$ in the bracketed term scales as $O(|\hat{V}_{\rm ext}|^{-1})$, and therefore $\tilde{I}^{\rm o}_{\rm MF} \sim -[2(4\chi\tilde{\gamma}\lambda_{\rm D}/R_{\rm P}+\tilde{\alpha}m)]^{-1}+\mathcal{O}(|\hat{V}_{\rm ext}|^{-1})$. Obtaining the next-order correction, we find an explicit relation between the mean-field current and the open fraction:

$$\tilde{I}^{\rm o}_{\rm MF} \;\sim\; -\frac{\chi+1}{\chi}\frac{\kappa_\infty}{1+2\hat{\alpha}m}\left(1+\frac{\hat{V}_{\rm res}}{|\hat{V}_{\rm ext}|}\right) \;, \tag{S.121}$$

where

$$\kappa_\infty \;=\; \frac{k_\infty}{k} \;=\; \frac{1}{8(\chi+1)\frac{\lambda_{\rm D}}{R_{\rm P}}\tilde{\gamma}} \;, \qquad \hat{V}_{\rm res} \;=\; \frac{\beta_{\rm ref}e\frac{\chi}{\chi+1}V_{\rm res}}{2} \;=\; \chi\left(1-\frac{\chi+1}{\chi}\frac{\kappa_\infty}{1-2\tilde{\gamma}}\right) \;, \tag{S.122}$$

$$\hat{\alpha} \;=\; \frac{\chi+1}{\chi}\kappa_\infty\tilde{\alpha} \;=\; \frac{\pi}{4(\chi+1)}\frac{R_{\rm P}L}{\ell^2} \;, \qquad \hat{\gamma} \;=\; \frac{\chi+1}{\chi}\kappa_\infty\tilde{\gamma} \;=\; \frac{R_{\rm P}}{8\chi\lambda_{\rm D}} \;. \tag{S.123}$$

In dimensional form, the current law becomes

$$I^{\rm o}_{\rm MF} \;\sim\; -k_\infty\mathrm{sgn}(V_{\rm ext})\frac{|V_{\rm ext}|+V_{\rm res}}{1+2\hat{\alpha}m} \;. \tag{S.124}$$

Here, $k_\infty$ and $V_{\rm res}$ are the effective channel conductance and residual voltage in the large-potential limit, as defined in Eq. (S.68). The conductance $k_\infty$ is determined by electrolyte transport properties since the concentration self-response, rather than the intrinsic channel resistance $k$, controls the overall current when the applied potential is large.

For the dielectric mismatch between the membrane and the electrolyte, $\chi \gg 1$, and the parameters in Table S1, the typical magnitudes of these dimensionless quantities are $\kappa_\infty \approx 4.6\times10^{-2}$, $\hat{\gamma}\approx 2.2\times10^{-3}$, and $\hat{V}_{\rm res}\approx 38$. Compared with the small-potential regime, both the apparent channel conductance and the channel-interaction strength are reduced by $\kappa_\infty/\kappa = \hat{\alpha}/\alpha \approx 7.5\times10^{-2}$, indicating that the concentration self-response suppresses the effective channel response under large applied potentials.

#### 3. Hamiltonian approximation

We aim to construct a Hamiltonian that describes the system dynamics in terms of $\{n_i\}$, both to draw parallels with the lattice/Ising model [37] and to obtain an expression for the mean open fraction as a function of the external voltage. To do so, consider that the force on each gating variable is given by

$$\frac{\partial U}{\partial \xi_i} \;=\; 4U^\ddagger(\xi_i^3-\xi_i) - Q(V_i - V_{\rm G}) \;, \tag{S.125}$$

from Eq. (S.70). Since $|Q(V_i-V_{\rm G})| \ll U^\ddagger$, we assume that the locations of the minima do not appreciably shift from $\xi_i=\pm1$. We may then compute the energy difference associated with flipping a channel from a closed state $n_i=0$ to an open state $n_i=1$ by integrating the above expression with respect to $\xi_i$ from $-1$ to $+1$:

$$\Delta U_i \;=\; -2Q\left[\frac{1}{2}\int_{-1}^{1} d\xi_i\; V_i(\xi_1,\ldots,\xi_i,\ldots,\xi_N) - V_{\rm G}\right] \;. \tag{S.126}$$

Under the separation of timescales argument, the only dependence of $V_i$ on $\{\xi_j\}$ arises through $n_i = \Theta(\xi_i)$. In that case,

$$\int_{-1}^{1} d\xi_i V_i(\{n_j\}) = \int_{-1}^{0} d\xi_i V_i(n_1, \ldots, n_{i-1}, 0, n_{i+1}, \ldots, n_N) + \int_{0}^{1} d\xi_i V_i(n_1, \ldots, n_{i-1}, 1, n_{i+1}, \ldots, n_N)$$
$$= V_i(n_1, \ldots, n_{i-1}, 0, n_{i+1}, \ldots, n_N) + V_i(n_1, \ldots, n_{i-1}, 1, n_{i+1}, \ldots, n_N) , \quad \text{(S.127)}$$

where the first term corresponds to the transmembrane potential when channel $i$ is closed and the second corresponds to that when channel $i$ is open, with all other channels held fixed. Under the mean-field current approximation, we have

$$V_i = \frac{\chi}{\chi+1} V_{\text{ext}} + \frac{I^{\text{o}}_{\text{MF}} n_i}{\pi g \lambda_{\text{D}}} + \sum_{\substack{j=1 \\ 0<r_{ij}<L}}^{N} \frac{\chi}{1+\chi} \frac{I^{\text{o}}_{\text{MF}} n_j}{\pi g r_{ij}} , \quad \text{(S.128)}$$

which gives

$$\Delta U_i = -2Q \left[ \frac{\chi}{\chi+1} V_{\text{ext}} - V_{\text{G}} + \frac{I^{\text{o}}_{\text{MF}}}{2\pi g \lambda_{\text{D}}} + \sum_{\substack{j=1 \\ 0<r_{ij}<L}}^{N} \frac{\chi}{1+\chi} \frac{I^{\text{o}}_{\text{MF}} n_j}{\pi g r_{ij}} \right] , \quad \text{(S.129)}$$

for the energy difference in terms of the mean-field open current $I^{\text{o}}_{\text{MF}}$.

We may then rewrite the energy change in Eq. (S.129) as

$$\Delta U_i = A + \sum_{\substack{j=1 \\ j\neq i}}^{N} B_{ij} n_j , \quad \text{(S.130)}$$

where

$$A = -2Q \left[ \frac{\chi}{\chi+1} V_{\text{ext}} - V_{\text{G}} + \frac{I^{\text{o}}_{\text{MF}}}{2\pi g \lambda_{\text{D}}} \right] , \qquad B_{ij} = \begin{cases} -2Q \dfrac{\chi}{1+\chi} \dfrac{I^{\text{o}}_{\text{MF}}}{\pi g r_{ij}} & 0 < r_{ij} < L \\ 0 & \text{otherwise} \end{cases} . \quad \text{(S.131)}$$

The coefficients $B_{ij}$ satisfy spatial invariance such that, for all $i$,

$$\sum_{\substack{j=1 \\ j\neq i}}^{N} B_{ij} = -2Q \sum_{\substack{j=1 \\ 0<r_{ij}<L}}^{N} \frac{\chi}{1+\chi} \frac{I^{\text{o}}_{\text{MF}}}{\pi g r_{ij}} = -\frac{4Q\tilde{\alpha} I^{\text{o}}_{\text{MF}}}{k} = B , \quad \text{(S.132)}$$

where $\tilde{\alpha}$ is the interaction term introduced previously. Furthermore, reciprocity holds, so that $B_{ij} = B_{ji}$ for all pairs $i$ and $j$.

We may now obtain a Hamiltonian that is consistent with the energy difference above for flipping the channel state $n_i$ from 0 to 1:

$$\mathcal{H}(\{n_i\}) = \sum_{i=1}^{N} A n_i + \frac{1}{2} \sum_{i=1}^{N} \sum_{\substack{j=1 \\ j\neq i}}^{N} B_{ij} n_i n_j , \quad \text{(S.133)}$$

where the factor of 1/2 accounts for double counting each interaction $B_{ij}$. Substituting the mean-field expansion $n_i = m + \delta n_i$, where $m$ is the mean open fraction, gives

$$\mathcal{H}(\{n_i\}) = -\frac{m^2 N}{2} B + (A + mB) \sum_{i=1}^{N} n_i + \mathcal{O}(\delta n_i \delta n_j) , \quad \text{(S.134)}$$

where we use the spatial invariance relationship. The mean-field approximation amounts to neglecting the quadratically coupled term in the Hamiltonian. Since each channel is independent in the mean-field Hamiltonian, the equilibrium distribution of $n_i$ is Boltzmann, giving the self-consistency equation

$$m = \frac{1}{1 + e^{\beta(A+mB)}} . \quad \text{(S.135)}$$

Equation (S.135) represents the mean open fraction of channels. Substituting the expressions for $A$ and $B$ gives

$$m = \left[1 + \exp\left(2\beta Q\left(V_{\mathrm{G}} - \frac{\chi}{\chi+1}V_{\mathrm{ext}} - \frac{I^{\mathrm{o}}_{\mathrm{MF}}}{k}\left[\tilde{\gamma} + 2\tilde{\alpha}m\right]\right)\right)\right]^{-1} . \tag{S.136}$$

This can be rewritten to obtain an explicit expression for the applied potential in terms of the open fraction as

$$V_{\mathrm{ext}} = \frac{\chi+1}{\chi}\left[V_{\mathrm{G}} - \frac{I^{\mathrm{o}}_{\mathrm{MF}}}{k}\left(\tilde{\gamma} + 2\tilde{\alpha}m\right) - \frac{k_{\mathrm{B}}T}{2Q}\ln\frac{1-m}{m}\right] , \tag{S.137}$$

which will be used in the subsequent analysis of the phase transition behavior.

#### 4. First-order transition and critical behavior

We now use the self-consistency conditions in Eqs. (S.111) and (S.137) to investigate the behavior of multiple channels. These two independent conditions relate the four variables $T$, $V_{\mathrm{ext}}$, $m$, and $I^{\mathrm{o}}_{\mathrm{MF}}$, leaving two degrees of freedom. Thus, specifying two variables, e.g. the control parameters $T$ and $V_{\mathrm{ext}}$, determines all the other variables, thereby defining the mean-field equations of state for the multiple-channel system. Explicit equations of state are generally not attainable due to the nonlinear and implicit nature of the current law in Eq. (S.111). In what follows, we thus focus on the limiting regimes of small and large applied potentials, which admit explicit equations of state, to analyze the phase transition and the critical behavior of the multiple-channel system.

##### (a). Small applied potentials

For small applied potentials, i.e. $|\hat{V}_{\mathrm{ext}}| \ll 1$, substituting the reduced current law in Eq. (S.119) into Eq. (S.137) gives the mean-field equation of state for $V_{\mathrm{ext}}$:

$$V_{\mathrm{ext}}(m,T) = \frac{\chi+1}{\chi}\left(\frac{1+2\alpha m}{1-\gamma}\left[V_{\mathrm{G}} - \frac{k_{\mathrm{B}}T}{2Q}\ln\frac{1-m}{m}\right]\right) , \tag{S.138}$$

where

$$\gamma = \frac{\chi+1}{\chi}\kappa\tilde{\gamma} = \frac{\chi+1}{\chi}\frac{k_{\mathrm{eff}}}{2\pi g\lambda_{\mathrm{D}}} . \tag{S.139}$$

As discussed in the main manuscript, $\gamma$ measures the effective channel conductance relative to the access-like conductance of the electrolyte near the channel pore. For larger $\gamma$, the local current perturbs the membrane potential more strongly in the direction that offsets the externally imposed potential. Thus, a stronger external potential is required to produce the same effective gating bias, as captured by the factor $(1-\gamma)^{-1}$ in Eq. (S.138). However, this local current correction cannot become arbitrarily large. Since the membrane capacitance is typically much smaller than the electrical double-layer capacitance, i.e., $\chi \approx 40$, the definition of $k_{\mathrm{eff}}$ in Eq. (S.64) imposes an upper bound, $\gamma \approx k_{\mathrm{eff}}/(2\pi g\lambda_{\mathrm{D}}) \lesssim 1/2$.

Equation (S.138) includes the mean-field interaction through the factor $1+2\alpha m$ and an entropy-like cost of maintaining a given open fraction, represented by the logarithmic term. Because of the collective interaction term, the equation of state can develop a segment with negative slope, $(\partial V_{\mathrm{ext}}/\partial m)_T < 0$, corresponding to unstable states. The two stable branches connected by this unstable segment are then separated by a discontinuous jump in the open fraction, indicating a first-order transition.

We first locate the critical point of this transition, at which the unstable segment shrinks to a single marginal state. There, the first and second derivatives of the equation of state vanish simultaneously:

$$\left(\frac{\partial V_{\mathrm{ext}}}{\partial m}\right)_T = \left(\frac{\partial^2 V_{\mathrm{ext}}}{\partial m^2}\right)_T = 0 . \tag{S.140}$$

Consider first the second-derivative condition. Since the contribution from $V_{\mathrm{G}}$ in Eq. (S.138) is linear in $m$, it does not contribute to the second derivative. Temperature then appears only as a common scale of the remaining terms and therefore does not affect the value of $m$ at which the collective interaction and entropic

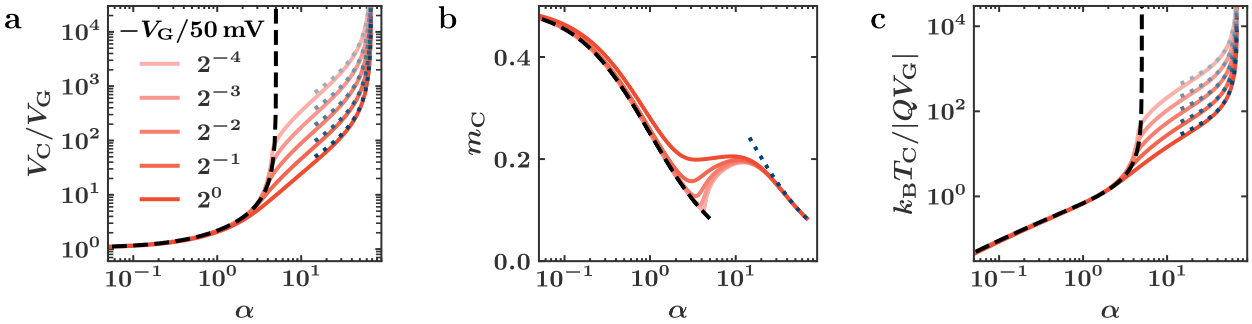

Figure S7: Mean-field critical behavior as a function of the channel interaction strength $\alpha$: (a) the critical applied voltage, (b) the critical open fraction, and (c) the critical temperature, using the parameters in Table S1. Solid red curves show numerical results obtained from the full nonlinear current relation in Eq. (S.114) for different gating voltages. Black dashed curves show the linearized small-potential result for $|\hat{V}_{\text{ext}}| \ll 1$ identified in Sec. V.4 (a), which captures the critical behavior accurately up to $\alpha \approx 5$, where the small-potential approximation becomes singular. Blue dotted curves show the large-potential result for $|\hat{V}_{\text{ext}}| \gg 1$ identified in Sec. V.4 (b), which captures the critical behavior accurately for $10 \lesssim \alpha \lesssim 67$. For $\alpha \gtrsim 67$, no critical point exists, and the first-order transition persists at any temperature.

effects balance to make the second derivative vanish. Consequently, the critical open fraction is determined by the second-derivative condition alone:

$$m_C = \frac{1}{2(1+\alpha)} , \tag{S.141}$$

which shows that the critical open fraction decreases with the channel interaction strength $\alpha$ and remains below the symmetric value of 1/2.

We next consider the first derivative,

$$\left(\frac{\partial V_{\text{ext}}}{\partial m}\right)_T = \frac{\chi+1}{\chi}\frac{2}{1-\gamma}\left[\alpha V_G + \frac{k_BT}{2Q}\left(\frac{1+2\alpha m}{2m(1-m)} - \alpha \ln\frac{1-m}{m}\right)\right] , \tag{S.142}$$

which diverges to positive infinity as $m \to 0$ or $m \to 1$. Thus, the minimum of the first derivative is located where the second derivative vanishes, i.e., $m = m_C$, and is given by

$$\left(\frac{\partial V_{\text{ext}}}{\partial m}\right)_T\Bigg|_{m=m_C} = \frac{\chi+1}{\chi}\frac{2}{1-\gamma}\left[\alpha V_G + \frac{k_BT}{2Q}\left(2\,(1+\alpha) - \alpha\ln(1+2\alpha)\right)\right] . \tag{S.143}$$

The unstable segment of the equation of state exists whenever this minimum value is negative. Thus, the criterion for a first-order transition is given by

$$V_G < -\frac{k_BT}{2Q}\left[\frac{2(1+\alpha)}{\alpha} - \ln(1+2\alpha)\right] , \tag{S.144}$$

which shows that the instability occurs more easily for smaller $V_G$. Note that the bracketed term in Eq. (S.144) becomes negative for $\alpha \gtrsim 5$. This implies two consequences of the linearized current law in Eq. (S.119). First, even channels with $V_G > 0$ can exhibit a first-order transition if the channel interaction is sufficiently strong. Second, for channels with $V_G < 0$, sufficiently strong channel interactions produce an instability at any temperature, so that no critical point exists.

However, the strong-interaction predictions for $\alpha \gtrsim 5$ should be interpreted as a limitation of the linearized current law in Eq. (S.119). The linearized form is valid only for the small-potential expansion of the channel current and therefore overestimates the collective effect in the strong-interaction regime. When the full current law in Eq. (S.114) is considered, the channel-interaction effect is regularized, and the above behaviors are no longer observed, as shown in Fig. S7.

A critical point exists if Eq. (S.143) can vanish, which determines the critical temperature:

$$\frac{k_B T_C}{-QV_G} = \frac{\alpha}{1+\alpha-\frac{1}{2}\alpha\ln(1+2\alpha)} \, . \tag{S.145}$$

For $\alpha \ll 1$, the critical point exists only for $V_G < 0$, and the critical temperature is proportional to $\alpha$ in this limit.

Substituting the critical open fraction and temperature into the equation of state, we obtain the critical applied voltage as

$$\frac{V_C}{V_G} = \frac{\chi+1}{\chi}\frac{1}{1-\gamma}\frac{1+2\alpha}{1+\alpha-\frac{1}{2}\alpha\ln(1+2\alpha)} \, . \tag{S.146}$$

We note that, whenever a critical point exists, the critical voltage is negative regardless of the gating voltage, implying that cooperative gating is essential for the first-order transition. Expanding about the critical point, we find mean-field Ising critical behavior, as expected from the mean-field formulation.

#### (b). Large applied potentials

We perform a similar analysis using the large-potential limiting behavior. We focus on $V_{ext} < 0$ to consider cooperative gating. Substituting the corresponding current relation in Eq. (S.124) into Eq. (S.137) gives the following equation of state:

$$V_{ext}(m,T) - V_{res} = \frac{\chi+1}{\chi}\frac{1+2\hat{\alpha}m}{1-\hat{\gamma}}\left(V_G - \frac{\chi}{\chi+1}V_{res} - \frac{k_B T}{2Q}\ln\frac{1-m}{m}\right) \, . \tag{S.147}$$

Except for the residual voltage offset $V_{res}$, which is retained from the subleading correction to the large-potential current law, Eq. (S.147) has the same structure as the small-potential equation of state in Eq. (S.138). Thus, the mechanism underlying the phase transition remains the same as the competition between the collective channel interactions and the entropy-like cost of maintaining the open fraction.

What changes in the large-potential limit is the appropriate apparent conductance: $k_\infty$ replaces its small-potential counterpart $k_{eff}$, yielding $\hat{\alpha}$ and $\hat{\gamma}$ instead of $\alpha$ and $\gamma$. From Eq. (S.123), the channel-interaction strength $\hat{\alpha}$ and the self-response strength $\hat{\gamma} \ll 1$ no longer depend on the transport parameters $g$ and $k$. Instead, they are determined by geometry, with both being proportional to the pore radius $R_P$, the length scale over which the concentration self-response develops. This reflects the fact that the concentration self-response controls the current in the large-potential limit.

We now locate the critical point using the conditions in Eq. (S.140), obtaining

$$m_C = \frac{1}{2(1+\hat{\alpha})} \, , \tag{S.148}$$

$$\frac{k_B T_C}{-Q\left(V_G - \frac{\chi}{\chi+1}V_{res}\right)} = \frac{\hat{\alpha}}{1+\hat{\alpha}-\frac{1}{2}\hat{\alpha}\ln(1+2\hat{\alpha})} \, , \tag{S.149}$$

$$\frac{V_C - V_{res}}{V_G - \frac{\chi}{\chi+1}V_{res}} = \frac{\chi+1}{\chi}\frac{1}{1-\hat{\gamma}}\frac{1+2\hat{\alpha}}{1+\hat{\alpha}-\frac{1}{2}\hat{\alpha}\ln(1+2\hat{\alpha})} \, , \tag{S.150}$$

The critical point in the large-potential limit has the same functional form as the corresponding result in the small-potential limit, Eqs. (S.141), (S.145), and (S.146), except that the gating voltage is shifted by the residual voltage. As in the small-potential limit, expanding the equation of state about the critical point gives mean-field Ising critical behavior.

For sufficiently strong channel interactions, where $\hat{\alpha} \gtrsim 5$, the critical point again vanishes for channels with $V_G < (\chi/(\chi+1))V_{res} \approx V_{res} \approx 2\,\mathrm{V}$, and the first-order transition persists at any temperature. This behavior is reflected in the divergence of the critical temperature and voltage at $\alpha \approx 67$, which corresponds to $\hat{\alpha} \approx 5$, in Fig. S7. Although channels with $V_G > (\chi/(\chi+1))V_{res} \approx V_{res} \approx 2\,\mathrm{V}$ may also admit a critical point for $\hat{\alpha} \gtrsim 5$ on the lower critical-temperature branch, such gating voltages are much larger than typical values and are therefore unlikely to be physiologically relevant.

##### (c). Phase diagrams

We now consider phase coexistence in the multiple-channel system. Under the mean-field Hamiltonian formulation in Sec. V.3, coexistence is determined by the Maxwell construction,

$$\int_{m_1}^{m_2} \left[V_{\text{ext}}(m,T) - V^*(T)\right] dm \;=\; 0 \;, \qquad \text{(S.151)}$$

where $m_1$ and $m_2$ are the coexisting open fractions and $V^*(T) = V_{\text{ext}}(m_1,T) = V_{\text{ext}}(m_2,T)$ is the external coexistence voltage. Although the binodal generally requires numerical evaluation, two limiting regimes can be analyzed explicitly: the low-temperature regime, $k_\text{B}T/|QV_\text{G}| \ll 1$, and the near-critical regime, $T \approx T_\text{C}$.

When $k_\text{B}T/|QV_\text{G}| \ll 1$, the roots of the spinodal condition $(\partial V_{\text{ext}}/\partial m)_T = 0$ approach $m = 0$ and $m = 1$. Since the binodal encloses the spinodal region, the coexisting open fractions also approach 0 and 1. The limiting behavior of the equations of state as $T \to 0$ shows that the coexistence voltage in this limit is governed by the gating voltage. For physiologically relevant channels, $|\hat{V}_\text{G}| \lesssim \mathcal{O}(1)$ and we thus consider the small-potential equation of state in Eq. (S.138) for the binodal condition in Eq. (S.151). This yields the leading order coexistence voltage as

$$V^*(T) \;\sim\; \frac{\chi+1}{\chi}\left[\frac{1+\alpha}{1-\gamma}V_\text{G} + \frac{\alpha}{1-\gamma}\frac{k_\text{B}T}{2Q}\right] \;. \qquad \text{(S.152)}$$

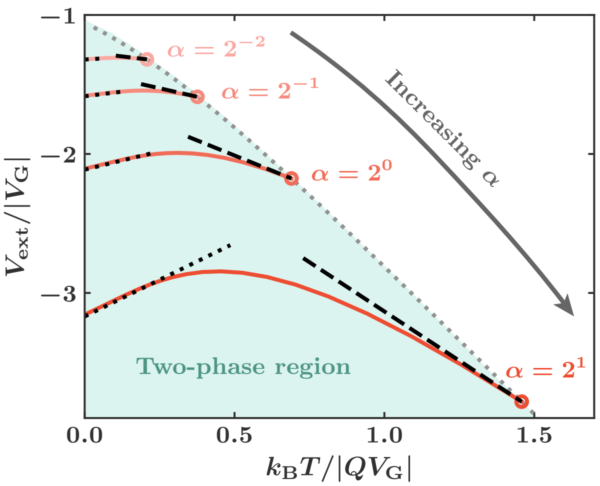

Figure S8: Phase diagram in the $V_{\text{ext}}$–$T$ plane for $V_\text{G} = -12.5\,\text{mV}$ and the parameters in Table S1. Red curves show coexistence curves for different channel interaction strengths $\alpha$, computed numerically using the full implicit equation of state determined by Eqs. (S.114) and (S.137). The gray dotted curve shows the critical line, along which $|V_\text{C}|$ and $T_\text{C}$ grow approximately proportionally to one another, as predicted by Eqs. (S.145) and (S.146). The black dotted lines show the low-temperature prediction for the coexistence voltage $V^*(T)$ from Eq. (S.152), and the black dashed lines show the near-critical prediction for $V^*(T)$ from Eq. (S.153).

This explains the weak temperature dependence of the coexistence potential for $k_\text{B}T/|QV_\text{G}| \ll 1$ in Fig. S8.

For the near-critical behavior, we expand the equation of state about the critical point and apply the binodal condition in Eq. (S.151). For $\alpha \lesssim 5$, the small-potential equation of state in Eq. (S.138) is valid and yields

$$V^*(T) \;=\; V_\text{C} + \frac{\chi+1}{\chi}\frac{k_\text{B}(T_\text{C}-T)}{2Q}\frac{1}{1-\gamma}\frac{(1+2\alpha)\log(1+2\alpha)}{1+\alpha} \;, \qquad \text{(S.153)}$$

where $V_\text{C}$ and $T_\text{C}$ are the critical voltage and temperature in Eqs. (S.146) and (S.145). For $\alpha \gtrsim 10$, the large-potential equation of state in Eq. (S.147) leads to the same form as Eq. (S.153), with $\gamma$ and $\alpha$ replaced by $\hat{\gamma}$ and $\hat{\alpha}$, respectively, and with $V_\text{C}$ and $T_\text{C}$ taken as the corresponding critical voltage and temperature in Eqs. (S.150) and (S.149). Equation (S.153) shows that the temperature dependence of the coexistence potential near the critical point increases with the channel interaction parameter $\alpha$, consistent with the numerically obtained binodal in Fig. S8.
